# Visceral organ morphogenesis via calcium-patterned muscle contractions

**DOI:** 10.1101/2021.11.07.467658

**Authors:** Noah P. Mitchell, Dillon Cislo, Suraj Shankar, Yuzheng Lin, Boris I. Shraiman, Sebastian J. Streichan

**Affiliations:** Kavli Institute for Theoretical Physics, University of California Santa Barbara, Santa Barbara, CA 93106, USA; Department of Physics, University of California Santa Barbara, Santa Barbara, CA 93106, USA; Department of Physics, Harvard University, Cambridge, MA 02138, USA; Biomolecular Science and Engineering, University of California Santa Barbara, Santa Barbara, CA 93106, USA

## Abstract

How organs achieve their final shape is a problem at the interface between physics and developmental biology. Organs often involve multiple interacting tissue layers that must be coordinated to orchestrate the complex shape changes of development. Intense study uncovered genetic, and physical ingredients driving the form of mono layer tissue. Yet, tracing dynamics across tissue layers, and scales – from cell to tissue, to entire organs – remains an outstanding challenge. Here, we study the midgut of *Drosophila* embryos as a model visceral organ, to reconstruct *in toto* the dynamics of multi-layer organ formation *in vivo*. Using light-sheet microscopy, genetics, computer vision, and tissue cartography, we extract individual tissue layers to map the time course of shape across scales from cells to organ. We identify the kinematic mechanism driving the shape change due to tissue layer interactions by linking out-of-plane motion to active contraction patterns, revealing a convergent extension process in which cells deform as they flow into deepening folds. Acute perturbations of contractility in the muscle layer using non-neuronal optogenetics reveals that these contraction patterns are due to muscle activity, which induces cell shape changes in the adjacent endoderm layer. This induction cascade relies on high frequency calcium pulses in the muscle layer, under the control of hox genes. Inhibition of targets of calcium involved in myosin phosphorylation abolishes constrictions. Our study of multi-layer organogenesis reveals how genetic patterning in one layer triggers a dynamic molecular mechanism to control a physical process in the adjacent layer, to orchestrate whole-organ shape change.

## I. INTRODUCTION

Visceral organ morphogenesis proceeds by the assembly of cells into tubes, which develop into complex shapes [1]. Through this process, genetic patterning instructs cellular behaviors, which in turn direct deformations in multiple interacting tissue layers to sculpt organ-scale shape. This motif arises, for instance, in the coiled chambers of the heart, contortions of the gut tube, and branching airways of the lung [2–4]. While studies of organ development in planar geometries imaged near the embryo surface or *ex vivo* have uncovered general principles governing cell and tissue behavior [5–10], mechanistic and dynamic understanding of visceral organ shape change has remained largely out of reach [11, 12]. Physical models inferred from static snapshots of organ morphology have proven useful in this regard [2, 13, 14]. The next step requires closing the gap between dynamics at the cellular and sub-cellular level with the dynamics of shape change at the organ scale.

Capturing these dynamics in a shape-shifting organ presents new challenges. Visceral organs exhibit both genetic and mechanical interactions between multiple tissue layers, supporting a wide palette of mechanistic complexity [15, 16]. Furthermore, since they develop deep inside embryos, capturing dynamics *in vivo* requires imaging methods that overcome image degradation due to scatter [17]. In addition, their complex, dynamic shapes hinder characterization even when imaging challenges are solved [18]. Here, we aim to tackle these challenges by studying the *Drosophila* midgut as a dynamic model system for *in toto* complex visceral organ shape change involving interacting tissue layers. The midgut – composed of muscle cells ensheathing an endodermal layer, linked by extracellular matrix (Fig. 1A) – offers a tractable system in which we can study the dynamics of morphogenesis and probe interactions across tissue layers. Its size and the molecular toolkit of the model system render the midgut ideal for light sheet microscopy [19], tissue cartography [20], and non-neuronal optogenetics [21]. The midgut displays gene expression patterns that are known to govern organ form and eventual cell fate [22–24]. For instance, hox genes expressed in the muscle layer are required for the midgut to form its four chambers, but the mechanism by which genetic expression patterns are translated into tissue deformation, and in turn to organ shape, remains unclear (Fig. 1B-C) [25–29]. Here we connect this genetic patterning to mechanical interactions between layers during development and track the kinematic mechanism linking mechanical action to organ shape transformations. We find that dynamic, high-frequency calcium pulses drive muscle contraction via the myosin light chain kinase, which efficiently induces bending in the bilayer tissue to sculpt stereotyped folds.

**FIG. 1.**
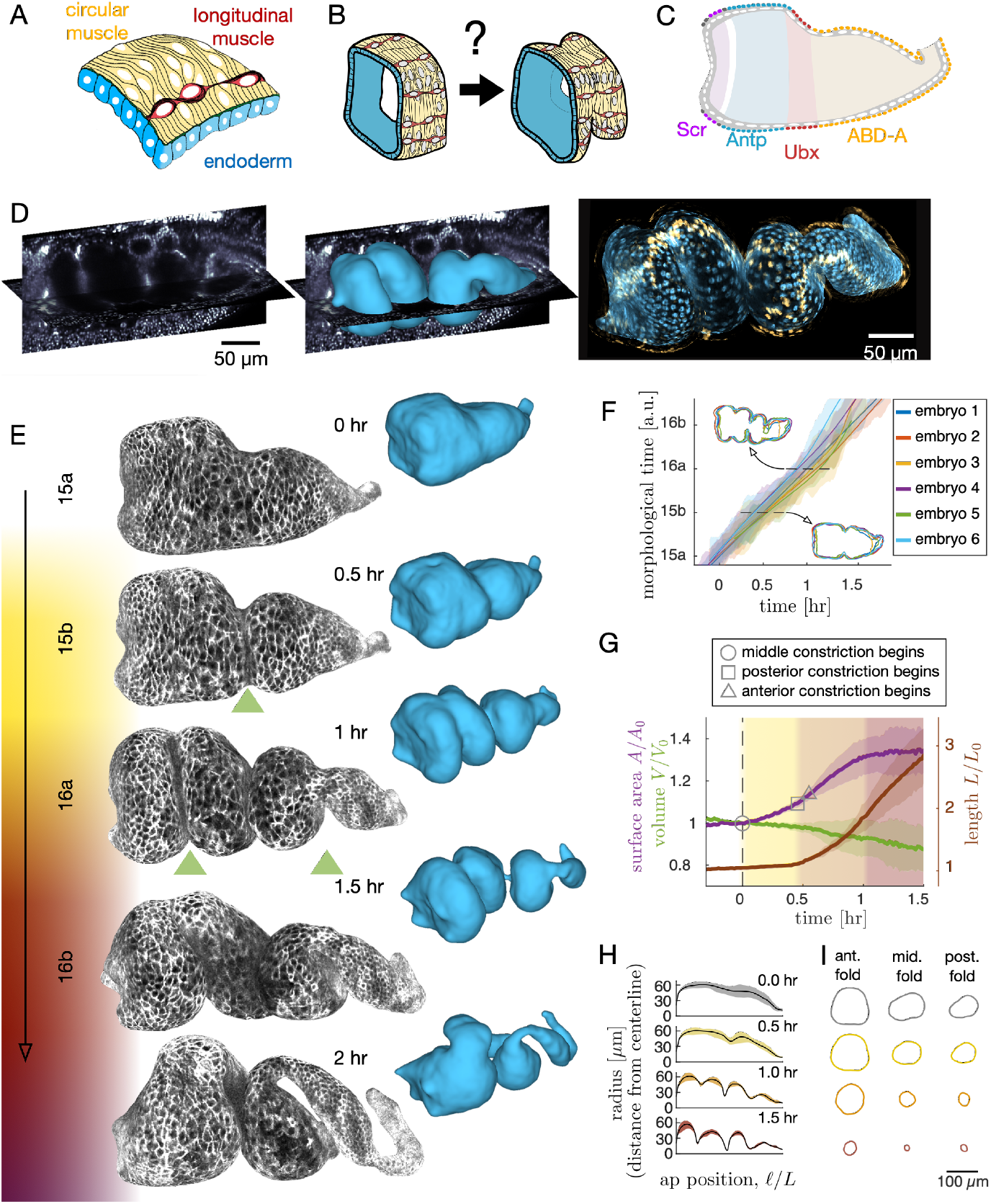
Gut morphogenesis is a stereotyped process of sequential constrictions in a bilayer tissue. *(A-B)* Muscle and endoderm layers compose the midgut and interact to generate 3D shape. *(C)* Genetic patterning of hox transcription factors that govern midgut morphogenesis appears in the circumferential muscles. *(D)* Automatic segmentation using morphological snakes level sets enables layer-specific imaging, here shown for muscle (yellow) and endoderm (blue) for a *w,Hand>GAL4;UAS-Hand:GFP;hist:GFP* embryo 90 minutes after folding onset. *(E)* Morphogenesis proceeds first with a constriction cleaving the gut into two chambers (stage 15b). Two more constrictions form a total of four chambers (16a) before the gut begins to coil (16b onward). Stages follow [30]. *(F)* Shape change is stereotyped, with shape transformations progressing at a reproducible rate, shown here for six morphologically-aligned datasets (see Supplementary Information). *(G)* Surface area of the apical surface increases during folding by ∼30%, but levels off by stage 16b. The enclosed volume decreases gradually throughout, while the effective length of the organ (computed via the length of a centerline) increases. *(H-I)* As folds deepen, constrictions adopt a circular shape. Snapshots of effective radius – measured as the distance from the centerline to circumferential hoops in the material coordinate frame, as a function of fractional position along the centerline – show tightening profiles near constrictions for a representative embryo. Colored bands bound the maximum and minimum effective radius. *(I)* Transverse profiles of each constriction for the same embryo used in (H) show circular constrictions in their respective dynamic cross-sectional planes.

## II. BILAYER MORPHOGENESIS PROCEEDS BY A REPRODUCIBLE PROGRAM OF SEQUENTIAL CONSTRICTIONS

The midgut is a closed tube by stage 15a of embryonic development (11 − 13 hours post fertilization) [31]. The organ first constricts halfway along its length, then constricts again to subdivide into four chambers. Within 75 − 90 minutes after the onset of the first fold, the constrictions are fully formed, and the organ adopts a contorted shape.

Quantitative characterization of these dynamics requires extraction of the full organ’s geometry. We overcome this challenge by combining *in toto* live imaging using MuVI SPIM with genetically-induced optical clearing and tissue specific markers (See Supplementary Information). To translate this volumetric data into geometric dynamics, we combine machine learning [32] with computer vision techniques [33, 34] and introduce an analysis package dubbed ‘TubULAR’ [35]. This captures all geometric information relating to shape change. We first extract the apical surface of the endoderm layer, and translate this surface through the thickness of the tissue to inspect specific layers, such as the yellow muscle layer in Fig. 1D.

From these extracted organ shapes, we find that gut morphogenesis is stereotyped and exhibits reproducible stages. By computationally aligning midgut shapes from different embryos, we find that the global shape varies little across embryos (Fig. 1F). The surface area grows by ∼30% during folding (stages 15a-16a) and remains constant by the time constrictions are fully formed (16b), despite continued shape change (Fig. 1G). The enclosed volume within the midgut decreases only gradually during this process, and the effective length of the organ – measured as the length of the centerline – triples by the time constrictions are fully formed (Fig. 1G). By parameterizing these surfaces with respect to their centerline, we see that constrictions become increasingly circular as folds deepen, despite the asymmetries of the overall shape (Fig. 1H-I, c.f. [27]).

## III. ENDODERMAL CELL SHAPE CHANGES LINK TO TISSUE SHAPE

How does this 3D shape change occur at the tissue and cellular scale? We first analyzed the endoderm layer. Inspection of these cells reveals striking anisotropic cell shapes before folding (Fig. 2A). In order to quantify cell shape on this dynamic surface, we cartographically project into the plane in a way that preserves geometric information (see Supplementary Information) [36]. The map to the plane also defines a global coordinate system in which we unambiguously define the anterior-posterior (AP) axis and circumferential (DV) axis for all time points, even when the organ exhibits deep folds and contortions (Supplementary Figure S5).

**FIG. 2.**
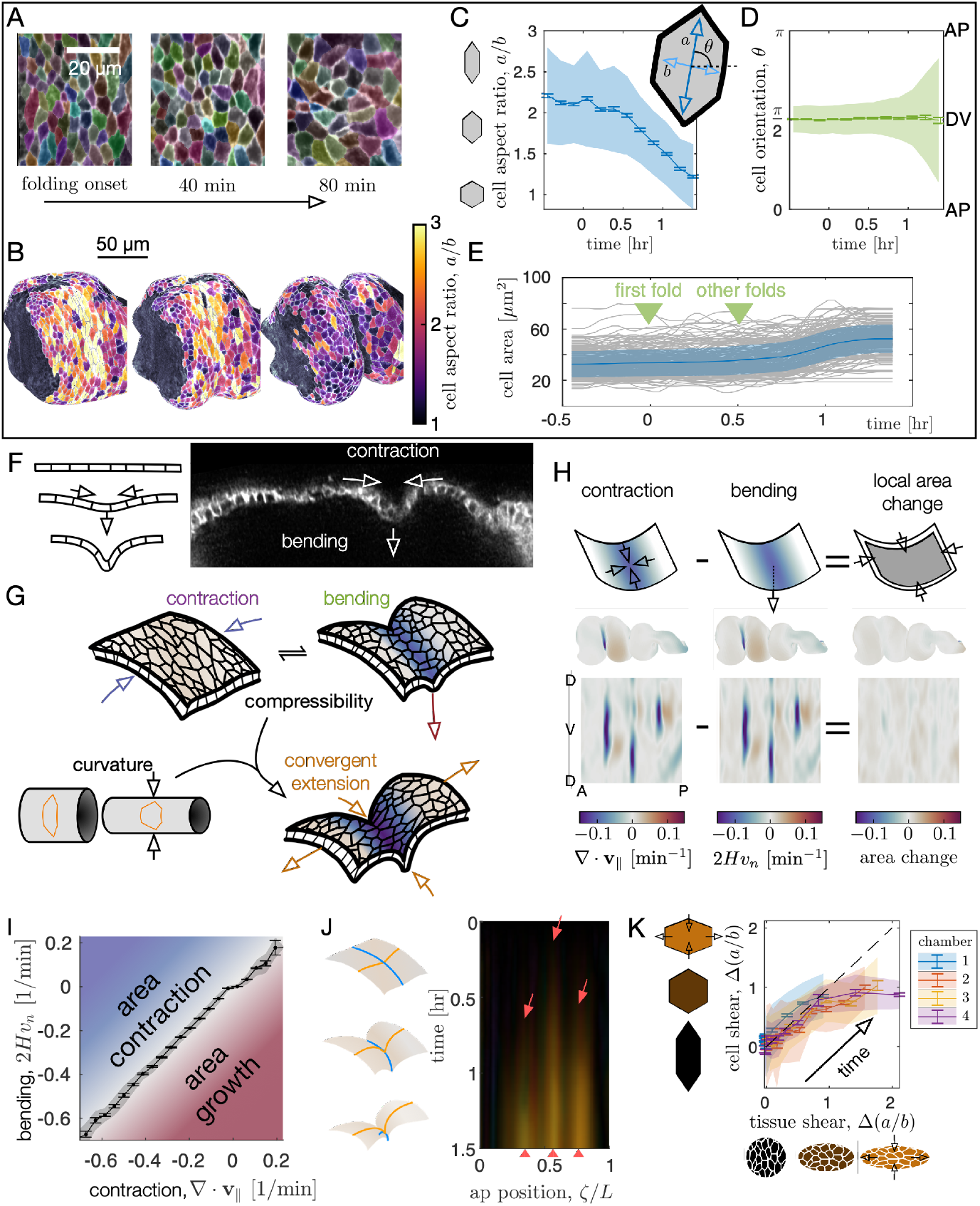
An interplay between tissue compressibility and organ geometry leads to convergent extension at both the cellular and tissue scale. *(A)* Cell segmentation in a computationally flattened coordinate patch shows endodermal cells are initially elongated along the circumferential direction but change their shape during organ folding. *(B-C)* Cell aspect ratios evolve from *a/b >* 2 to *a/b* ≈ 1 during folding, shown in 3D for cells near the anterior fold. Colored bands denote area-weighted standard deviations for 600-1300 segmented cells per timepoint, and tick marks denote standard error on the mean. *(D)* As cells change their aspect ratio, their orientations do not rotate. *(E)* Single-cell tracking shows gentle increase of cell areas through violent folding events, suggesting that cell area changes do not drive organ shape change. *(F)* Constrictions display contractile in-plane velocity and out-of-plane bending. The degree of compressibility in the tissue – here nearly incompressible – links in-plane contraction to out-of-plane bending. *(G)* Convergent extension via circumferential constriction – in which tissue converges along the DV direction and extends along the bending AP profile – follows as a geometric consequence of constricting the tubular midgut without local area change. *(H)* Global measurements of tissue deformation reveal tight coupling between dilatational flow and bending (∇· **v**_∥_≈ 2*Hv*_*n*_), with only a weak local area change. *(I)* Divergence and bending are correlated at the 97% level, signalling nearly incompressible behavior (*N* = 3 embryos, with kinetics sampled in 320 tissue patches per timepoint for 0 *< t <* 90 min). Gray band denotes standard deviation and ticks denote standard error on the mean for each bin. *(J)* Tissue-scale shear accumulates near each fold, shown as a kymograph in the material coordinate frame averaged along the circumferential axis. As time increases (downward), orange streaks reflect area-preserving convergence of tissue patches along the organ circumference and extension along the folding AP axis near constrictions (red arrows). *(K)* Tissue-scale shear is mirrored in cellular shape change in all chambers. The change in cell aspect ratio is plotted against the change in tissue patch aspect ratio when advected under the mean tissue velocity. Each data point is an average over hundreds of cells in a given chamber, with standard deviations denoted by colored bands and standard error by tick marks.

By segmenting cells in 2D and mapping their contours back into 3D, we find that endodermal cells are strongly anisotropic, with an average aspect ratio *a/b >* 2, and are globally aligned along the circumferential axis (Fig. 2A-C). The endodermal cells lose their initial anisotropy as the folds develop, and even become elongated along the AP axis in some regions by the end of folding (see Supplementary Information). Measurement of endodermal cell orientations reveal that this effect is not due to rotations (Fig. 2D). Despite the large changes in aspect ratio, cell areas in the endoderm layer change only gradually during folding (Fig. 2E), so cells converge along the circumferential axis while extending along the organ’s longitudinal axis. On large scale, this cell deformation would collectively generate tissue movement corresponding to convergent extension. We would intuitively expect these cell shape changes to accompany narrowing and lengthening of a tube constricting uniformly. However, constrictions are strongly localized along the AP axis, making the relationship between cell shape change and 3D tissue shape unclear *a priori* (Fig. 2F-G).

To understand the kinematic mechanism underlying organ shape, we must bridge spatial scales from cell deformation to meso-scale tissue deformation. To this end, we extract whole-organ tissue deformation patterns and find strong in-plane contractile flows at folds, where the out-of-plane bending is large (Fig. 2F-H). How do these motions couple to generate 3D form?

In 3D, tissue can stretch, bend, and shear. Unlike in flat or confined geometries, here cells need not change their area in response to in-plane contractile flows: the tissue can instead bend into the out-of-plane direction to preserve area (Fig. 2F-G) [37]. We hypothesized that the tissue could behave as an incompressible medium, which would imply that in-plane contraction and out-of-plane bending are tightly linked. Remarkably, we find that contraction patterns almost entirely account for bending patterns, with only a small change in local tissue areas (Fig. 2H). As shown in Fig. 2I, the bending and contraction terms match with 97% correlation, leaving a residual in-plane growth residue in the lobes of the gut chambers at the level of ≲ 1% per minute. This slow area growth accounts for the cellular surface area growth noted in Fig. 1G.

As a consequence of this nearly area incompressible behavior, the fact that the organ is curved in the circumferential direction mandates that inward motion of constrictions must generate convergent extension (Fig. 2G). We dub this kinematic mechanism ‘convergent extension via circumferential constriction:’ convergence along the circumferential direction is not accompanied by tissue velocity along that direction, but instead in the out-of-plane direction. The tissue converges circumferentially and extends along its longitudinal (AP) axis by folding, despite the fact that cells flow into the fold from both the anterior and the posterior (Fig. 2J). This behavior is only possible because of the localized tissue’s curvature and out-of-plane motion (see Supplementary Information). Fig. 2J shows a kymograph of the accumulated shear deformation in a representative embryo, showing deformation is strongest near each constriction. Though the shape of the organ becomes increasingly complex as morphogenesis proceeds, shear deformations remain globally aligned: the tissue converges (extends) along the organ’s intrinsic DV (AP) axis, denoted by the nearly uniform color of streaks in Fig. 2J.

Finally, we find this tissue shear deformation reflects our previous measurement of cell shape change (Fig. 2K). The correspondence between cellular shear and tissue shear indicates that local cell shape changes primarily mediate meso-scale convergent extension. In short, we established a link from endodermal cell shear to tissue-scale folding in which strong, localized contractile flows are translated directly to out-of-plane bending, resulting in convergent extension via circumferential constriction. What cellular process drives these strong localized contractions at the folds?

## IV. MUSCLE CONTRACTIONS DRIVE CELL AND TISSUE SHAPE CHANGE

It is known that embryos with either deleted muscle or endoderm structure fail to fold normally [24, 38–40], and removal of integrins linking the two layers likewise results in uncontrolled shapes [29]. This suggests that gut morphogenesis requires an interaction between muscle and endodermal layers. At the same time, hox genes expressed only in the muscle layer are known to be upstream of the constriction process, such that removing particular hox genes removes the associated folds (Fig. 3A-B) [41]. In particular, *Antp* mutants lack the anterior fold lying near the center of the *Antp* domain, while *Ubx* mutants lack the middle fold lying at the posterior edge of the *Ubx* domain (Fig. 3B). Is the interaction required for constrictions genetic in nature – with muscle cells relaying instructions to the endoderm [22] – or mechanical – with active stresses in the muscle layer inducing endodermal cell shape change [27]? To clarify the relationship between layers during constriction dynamics, we first measured relative motion of the muscle layer against the endoderm. By tracking both circumferential muscle nuclei and endoderm nuclei in the same embryo, we find that these two layers move together, with an resulting displacement between initially close nuclei of ∼5*µ*m per hour (Fig. 3C and Supplementary Information). This result is consistent with the notion that the two layers are tightly tethered by the integrins and extracellular matrix binding the heterologous layers [29].

**FIG. 3.**
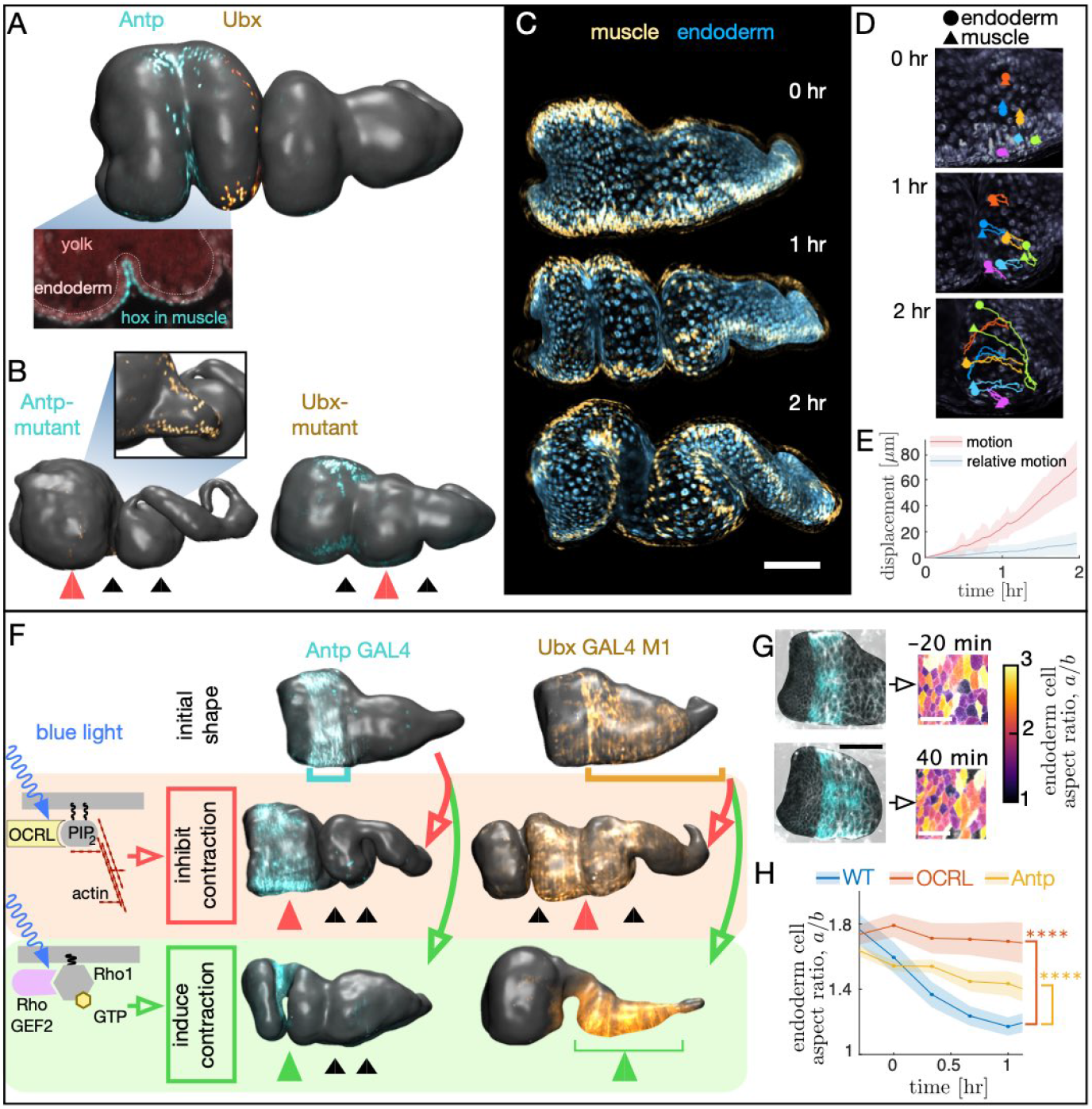
Muscle contraction changes organ shape and endoderm cell shapes via mechanical coupling between layers. *(A)* Hox genes *Antp* and *Ubx* are expressed in the circumferential muscle in regions spanning the anterior fold and anterior to the middle fold, respectively. *(B)* Hox genes control organ shape: *Antp* and *Ubx* mutants lack anterior and middle constrictions, respectively. *(C-D)* Muscle and endoderm layers move together. By computationally extracting both muscle and endoderm layers in an embryo expressing both fluorescent circumferential muscle (*Hand-GAL4* ; *UAS-Hand:GFP*) and endoderm (*hist:GFP*), we track relative motion of initially close muscle-endoderm nuclei pairs. *(E)* Muscle-endoderm nuclei pairs show modest relative motion compared to the integrated motion of the tissue (*N* = 81 pairs, colored bands denote standard deviations). *(F)* Optogenetic inhibition of contractility via *OCRL* mimics hox mutant behaviors (*N* = 11), and stimulation via *UAS-RhoGEF2* in the muscle layer drives ectopic folding (*N* = 5). *(G)* Inhibiting muscle contraction prevents endoderm cell shape change, shown for snapshots before and after the anterior constriction would normally form. *(H)* Measurements of endodermal cell anisotropy over time confirm that mechanical inhibition in the muscle reduces cell shape change in the endoderm (*p* = 1 × 10^−22^). *Antp* mutants also exhibit reduced endoderm cell shape change, consistent with *Antp* regulating muscle contraction (*p* = 4×10^−9^). Each datapoint is the weighted average of multiple adjacent timepoints and 2-3 embryos with at least 30 cells per timepoint segmented in each embryo. Colored bands denote standard error on the mean.

Based on this tight coupling, we hypothesized that muscle mechanically induces shape change in the tethered endoderm. To test this hypothesis, we inhibited contractility of the muscle layer using an optogenetic construct under the control of hox-gene-based GAL4 drivers. Under these optogenetic conditions, the genetic patterning remained intact, but the mechanical behavior of the muscle was altered. Driving *OCRL* with *Antp-GAL4* under continuous activation of blue light reliably prevented anterior folding (Fig. 3F). Likewise, driving *OCRL* under continuous blue light activation in muscle regions posterior to the anterior fold using *Ubx-GAL4 M1* inhibited constriction dynamics, leading to only a transient and partial middle fold and a delayed posterior fold. We note that *Ubx-GAL4 M1* embryos express *Ubx* in a larger domain than the endogenous WT *Ubx* domain due to differences in its regulation, but *Ubx-GAL4 M1* embryos nonetheless execute all three folds in the absence of *UAS-OCRL* under similar imaging conditions. Inhibiting contraction in selected regions therefore mimics the genetic mutants known to remove folds.

Given that muscle contractility is required, we asked if optogenetically inducing muscle contraction is sufficient to induce constrictions. Indeed, optogenetic activation using the *CIBN UAS-RhoGEF2* system in the *Antp* region generates an anterior fold on demand on the timescale of a few minutes, even if induced long before the constriction would normally begin (Fig. 3F). Similarly, activation of the *Ubx-GAL4 M1* domain, which is far more spatially extended along the AP axis, results in a nearly uniform constriction that dramatically alters the shape of the organ, forcing the yolk to flow into the anterior chamber. Additional optogenetic experiments inhibiting contractility of all muscles likewise led to folding defects (N=13). We conclude that muscle contractility is necessary for constrictions and inducing contraction and the associated downstream behaviors is sufficient to generate folds.

We then asked how these macro-scale perturbations on organ shape are linked to cell shapes in the endoderm. In contrast with the wild-type, endodermal cell shape changes are significantly reduced under optogenetic inhibition of muscle contractility. As shown in Fig. 3G-H, cell segmentation of the endoderm during optogenetic inhibition of muscle contraction in the *Antp* domain reveals nearly constant aspect ratios: the endoderm cells near the *Antp* domain undergo reduced convergent extension when muscle contraction is locally disrupted (single-sided z-test: *p* = 1 × 10^−6^ for difference after 1 hr, *p* = 1 × 10^−22^ for sustained difference between curves, see Supplementary Information). We also observe analogous reduction of endodermal cell shape change in *Antp* mutants, which lack anterior folds (single-sided z-test: *p* = 7 × 10^−3^ for difference after 1 hr, *p* = 4 × 10^−9^ for sustained difference between curves). Thus, the endodermal program of convergent extension is induced by mechanical interaction with the contracting muscle layer.

## V. CALCIUM PULSES SPATIOTEMPORALLY PATTERN MUSCLE CONTRACTILITY

What mechanism triggers muscle contractions, allowing such sharp folds to arise [42]? Recent studies have shown that calcium signaling triggers muscle contractions in a wide range of contexts [43]. If hox genes use calcium signaling to pattern muscle contraction in the midgut, we would predict that calcium pulses appear near localized constrictions. Furthermore, hox gene mutants lacking folds would not exhibit localized calcium pulses, and inhibition of the cell biological mechanism translating calcium into mechanical contraction should likewise inhibit constrictions.

To tested for a link from hox genes to organ shape through this mechanism, we fist imaged a fluorescent probe of calcium dynamics (*GCaMP6s*) in the muscle layer. As shown in Fig. 4A-F, transient calcium pulses appear in the muscle layer in regions localized near all three midgut constrictions. Additionally, these calcium pulses are patterned in time, appearing only at the onset of constriction for each fold (see Supplementary Information).

**FIG. 4.**
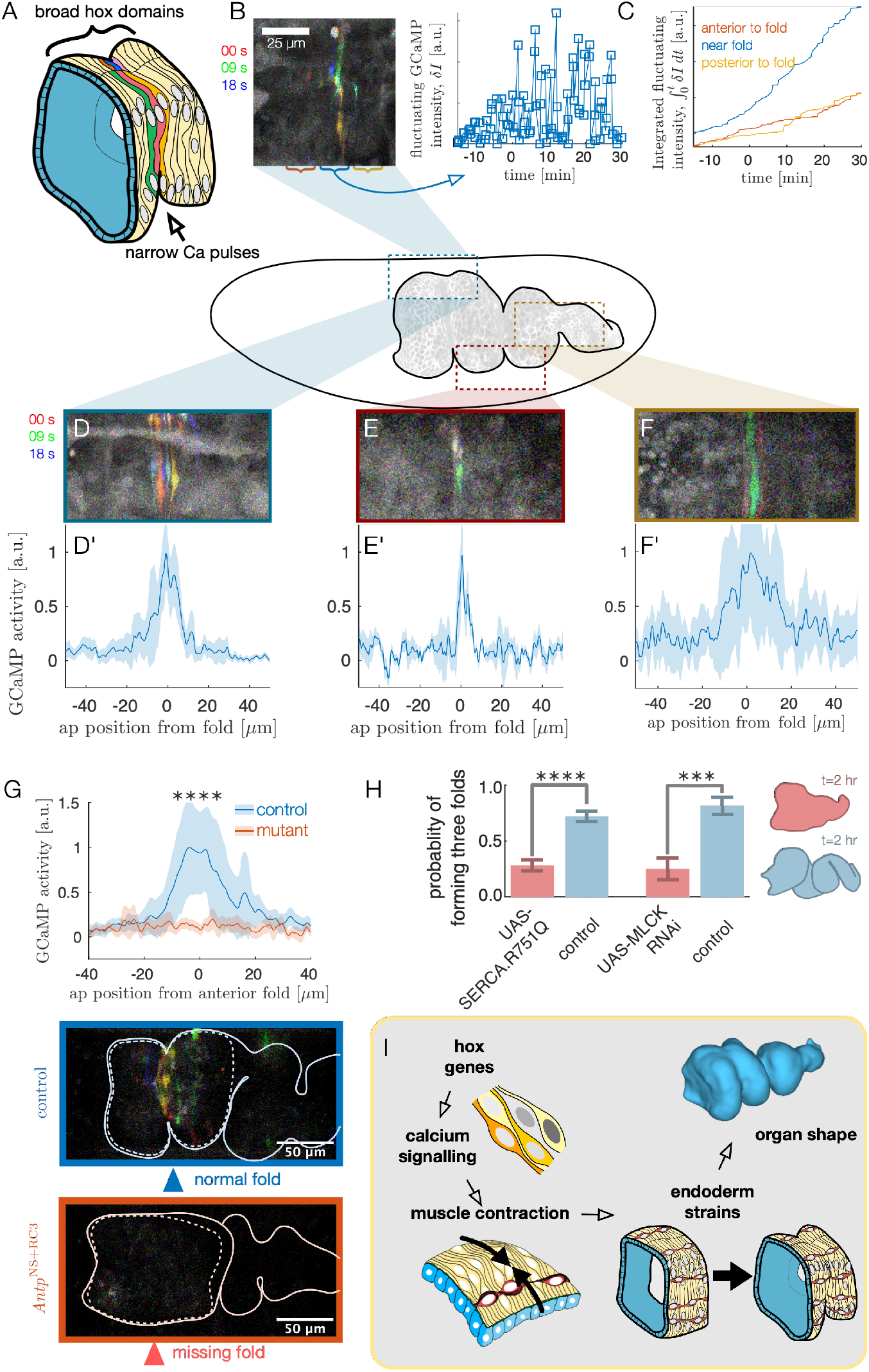
High frequency pulsatile calcium dynamics mediate muscle contraction, establishing a multi-scale link from hox genes to shape through tissue mechanics. *(A)* Dynamic calcium pulses appear near the anterior fold, localized to a region more narrow than the *Antp* domain. *(B)* Transient pulses in *GCaMP6s* intensity occur on the timescale of seconds and increase in amplitude when folding begins (*t* = 0). Red, green, and blue channels of images represent maximum intensity projections of confocal *z*-stacks separated in time by 9 seconds. *(C)* Integrated transient pulses for the same embryo as in (B) shows calcium pulses are localized near the fold: *GCaMP6s* signals 20*µ*m in front (red) or behind the fold (yellow) are less intense. *(D-F)* Snapshots of *GCaMP6s* fluorescence in muscle cells demonstrate calcium activity near constrictions. Here, we used *Mef2-GAL4* as a driver for characterizing anterior and middle constrictions, and *48Y GAL4* provided similar results. We used *48Y GAL4* for characterizing *GCaMP6s* fluorescence near the posterior constriction, since many fluorescent somatic muscles occlude the line of sight for the posterior fold under *Mef2-GAL4*. *(D’-F’)* Average fluorescent activity during the first 15 minutes of folding show localized signatures, with particularly sharp peaks in the middle and anterior constrictions. *N* = 5, *N* = 2, and *N* = 7 for anterior, middle, and posterior folds, respectively. *(G)* In *Antp* mutants, *GCaMP6s* fluorescence is significantly reduced (*p* = 2 × 10^−8^) and is not localized in space. Snapshots of *GCaMP6s* expression 28 minutes after posterior fold onset show almost no activity in the anterior region compared to the control. *(H)* Disruption of calcium regulation in muscle cells inhibits constrictions. The probability of forming three folds is reduced under heat-shock induced expression of the dominant negative mutant allele *SERCA*.*R751Q* with a muscle-specific driver *Mef2-GAL4* (*p* = 7 × 10^−9^), and is likewise reduced under RNA interference of MLCK driven by *tub67-GAL4; tub16-GAL4* (*p* = 2 × 10^−4^). *(I)* Altogether, we infer that hox genes are upstream of patterned calcium pulses, which generate muscle contraction that is mechanically coupled to the endoderm, driving tissue strains and ultimately organ shape.

To test whether hox genes pattern shape change through calcium dynamics, we measured *GCaMP6s* activity in flies mutant for *Antp* that lack an anterior constriction. As shown in Fig. 4G, we found that calcium activity was almost entirely absent during stages 15-15b. Calcium activity is strongly reduced at the location of the missing anterior fold (single-sided z-test: *p* = 2 × 10^−8^) and subsequent calcium pulses are repressed within the vicinity of the region for the hour after folding would normally initiate (single-sided z-test: *p* = 1 × 10^−13^ within 50 *µ*m of the anterior fold location). The hox gene *Antp* is therefore upstream of dynamic calcium pulses.

Importantly, we also find that in wild-type embryos, knock-downs of calcium signaling remove folds. Under suitable release into the cytosol, calcium is known to trigger muscle contraction by binding to calmodulin, which in turn binds to myosin light chain kinase (MLCK) to trigger myosin light chain phosphorylation [44], and cytoplasmic calcium is transported from the cytosol into the sarcoplasmic reticulum for storage under regulation of SERCA [43]. We find that driving expression of a dominant negative form of SERCA via heatshock beginning at stages 13-15a suppressed midgut constrictions (*p* = 7 × 10^−9^, Fig. 4H). Separately, interrupting the production of MLCK in the muscle via RNA interference (RNAi) demonstrates a similar reduction in folding behavior (*p* = 2 × 10^−4^, Fig. 4H). From this we infer that spatially localized calcium dynamics – under the control of hox gene patterning – triggers MLCK signaling leading to muscle contractions.

## VI. DISCUSSION

Here we studied morphogenesis of an organ in which heterologous tissue layers generate complex shape transformations. We found that convergent extension and bending in the endodermal layer is triggered by mechanical interaction with muscle contractility, and patterns of calcium signalling regulate contractility in muscle cells according to hox-specified information (Fig. 4I).

Though correspondences between hox genes and cell fates have been established for decades [45], understanding the physical processes driven by hox genes remains an active area of research. Here we demonstrate a link from genes to tissue morphodynamics through active forces mediated by calcium signaling. High-frequency calcium pulses pattern muscle contraction in circumferential (DV) stripes positioned along the longitudinal (AP) axis of the organ, leveraging the anisotropy of initial cell shapes to generate anisotropic contractions. These contractions in turn induce tissue deformation through a motif of convergent extension that relies on circumferential constrictions and the nearly incompressible behavior of the tissue. As endodermal cells flow into the deepening folds, cell shapes converge along the circumferential direction while extending along the folding AP axis to accommodate the extreme changes in geometry without changing area. This induction cascade integrates high frequency calcium pulses to advance reproducible morphogenesis of complex 3D shape.

At the tissue level, a remaining question is to what extent endodermal cells actively respond to muscle contraction, rather than passively deforming. For instance, there could be a mechanical signaling pathway provoking behavior in endodermal cells. At the cellular level, a remaining question is how the midgut selects precise positions for localized calcium activity despite broad hox gene domains. In the anterior fold, for example, forms near the center of the *Antp* domain (see also Supplemental Material). Do cells sense subtle gradients of *Antp*, or does more refined patterning – either downstream or independent of hox gene expression – specify this location [42]? One available avenue for the latter possibility is that hox genes govern the formation of anatomical structures that could allow external signals from the soma to trigger calcium pulses in specific locations. A fold-specific candidate is the alary-related thoracic muscle connected to the dorsal side of anterior midgut [46]. The timing mechanism instructing cells to begin contraction additionally remains unclear [38]. Observing midgut development in the absence of the soma presents a future challenge that could address these questions.

Our observations relate more broadly to the ability of ion signaling to govern morphogenetic processes [47, 48]. Calcium dynamics have roles in early developmental stages of plants, vertebrates, and invertebrates alike [49–53], including regulating visceral organ growth [54]. In vertebrates, peristaltic waves of calcium pulses have recently been shown to influence the organization of muscle fibers in the midgut [16], and calcium-mediated signalling determines cell fates in heart valves [55]. Our findings demonstrate that patterned calcium pulses control 3D shape through a mechanical cascade across tissue layers.

## VII. ACKNOWLEDGEMENTS

This research was supported by NIH Grant No. R35 GM138203 and NIH Grant No. R00 HD088708, and was supported in part by the National Science Foundation Grant No. NSF PHY-1748958. We thank members of the Streichan and Shraiman labs, Eric Wieschaus, and Zvonimir Dogic for valuable insights, discussions, and suggestions. Isaac Breinyn aided in early exploration of the system and quantification of midgut morphology. Isaac Breinyn, Sophie Streichan, and Matt Lefebvre aided in handling several stocks, crosses, and reagents. We acknowledge Ben Lopez in the NRI-MCDB Microscopy Facility for support and maintenance of the Resonant Scanning Confocal supported by the NSF MRI grant DBI-1625770. The authors acknowledge the use of the Microfluidics Laboratory (Innovation Workshop) within the California NanoSystems Institute, supported by the University of California, Santa Barbara and the University of California, Office of the President. NPM acknowledges support from the Helen Hay Whitney Foundation. SS acknowledges support from the Harvard Society of Fellows.

