## Supplemental information for "Visceral organ morphogenesis via calcium-patterned muscle contractions"

### I. STOCKS AND REAGENTS

For this work, we used the following stocks based on Bloomington Drosophila Stock Center stocks:

*48Y-GAL4; klar*,

*w; UAS-CAAX::mCherry; +*,

*w; UAS-histone::RFP*,

*w; UAS-LifeAct::Ruby; +*,

*w; +; UAS-LifeAct::RFP*,

*UAS-MLCK RNAi TRiP VALIUM 20*,

*y,w,Antp-GAL4; +; klar*,

*w; sqh GFP; +*,

*w; +; Mef2-GAL4*

*w[*]; I-76-D,* $Ubx^{9.22}$ $e^1$*/TM6B, Tb*

$Df(3R)Antp^{NS+RC3}$.

Additional stocks used were

*w[*]; UASp>CIBN::pmGFP; UASp>mCherry::CRY2-OCRL* (a gift from Stefano de Renzis)

*w[*]; UASp>CIBN::pmGFP; UASp>RhoGEF2-CRY2::mCherry* (a gift from Stefano de Renzis)

*Hand-GFP; 4xUAS-Hand; +* (a gift from Zhe Han),

*w; +; Miple2-GAL4 #2* (a gift from Ruth Palmer) [1],

*w; +; Miple2-GAL4 #4* (a gift from Ruth Palmer),

*w; +; Ubx-GAL4 M1* (a gift from Ellie Heckscher) [2, 3],

*w; +; Ubx-GAL4 M3* (a gift from Ellie Heckscher) [2, 3],

*w; tub67>CAAX mCherry;+*,

*w; tub67-GAL4;tub15-GAL4*,

*w; +; tub67>CAAX mCherry*,

*w; +;* $Gap^{43}$ *mCherry.*

We used following antibodies for staining: Ubx FP3.38 (diluted 1:10, Developmental Studies Hybridoma Bank), Antp 8C11 (diluted 1:25, Developmental Studies Hybridoma Bank), anti-ABDA C11 (dilued 1:50, Santa Cruz Biotechnology).

---

*
†

### II. MICROSCOPY

For light sheet imaging of live and fixed embryos, we used a custom multi-view selective plane illumination microscope drawn schematically in Fig. S1A) [4]. As in [5], two illumination objectives (CFI Plan Fluor 10x, NA 0.3) focus a laser beam on a given embryo embedded in an agar column. The illumination path features discrete laser lines as sources (OBIS 488nm, 561nm, 633nm), a tube lens (200 mm, Nikon Instruments Inc.), and a scan lens (S4LFT0061/065, Sill optics GmbH and Co. KG). Fluorescence is captured by two detection objectives aligned in the orthogonal direction (App LWD 5x, NA 1.1, Nikon Instruments Inc.). Each embryo is mounted in an agar column with its anterior-posterior axis along the long axis of the column, and the column is oriented transverse to both the axis formed by the illumination objectives and the axis formed by the detection objectives (Fig. S1). The illuminating beams are translated up and down via mirror galvanometers (6215 hr, Cambridge Technology Inc.), sweeping through $y$ in tandem with the CMOS pixel row readout of two cameras (not shown) to form an image in each $xy$ plane (black squares in Fig. S1B). By translating the embryo through a series of such planes spaced $1.2-1.4$ $\mu$m apart using Optical section employed a translation stage (Physik Instrumente GmbH and Co. KG, P-629.1CD with E-753 controller), a data volume is formed (stack of black frames in Fig. S1B). By rotating the embryo through two or more angles (U-628.03 with C-867 controller, 0 and 90 degree views are shown in blue and yellow in Fig. S1B), a multi-view data volume is acquired.

Subsequent registration, deconvolution, and fusion using the methods presented in [6] results in a single, deconvolved data volume per timepoint with isotropic resolution (0.018 $\mu\text{m}^3$ per voxel). For most experiments in this work, we acquire one volume per minute. The optimal number of deconvolution iterations varied between 8 and 20 for different fluorescent reporters. We used 6-8 views for most datasets.

To peer deep inside the developing embryo, we leverage the UAS-GAL4 system [7] to express fluorescent proteins in gut-specific tissues and use embryos with the *klarsicht* mutation, which enables deep tissue imaging without altering morphogenesis. Fig. S2 shows a maximum intensity projection of the left lateral half data volume for a deconvolved nuclear marker in the midgut as a complement to Figure 1 of the main text. In Fig. S2, non-midgut tissue expressing *48Y GAL4* are largely masked out, but some cells outside the midgut that express *48Y GAL4* are visible on the dorsal side.

We used confocal microscopy (Leica SP8) for more detailed characterization of calcium dynamics, live imaging of *Ubx* and *Antp* mutants, and supplementary optogenetic experiments.

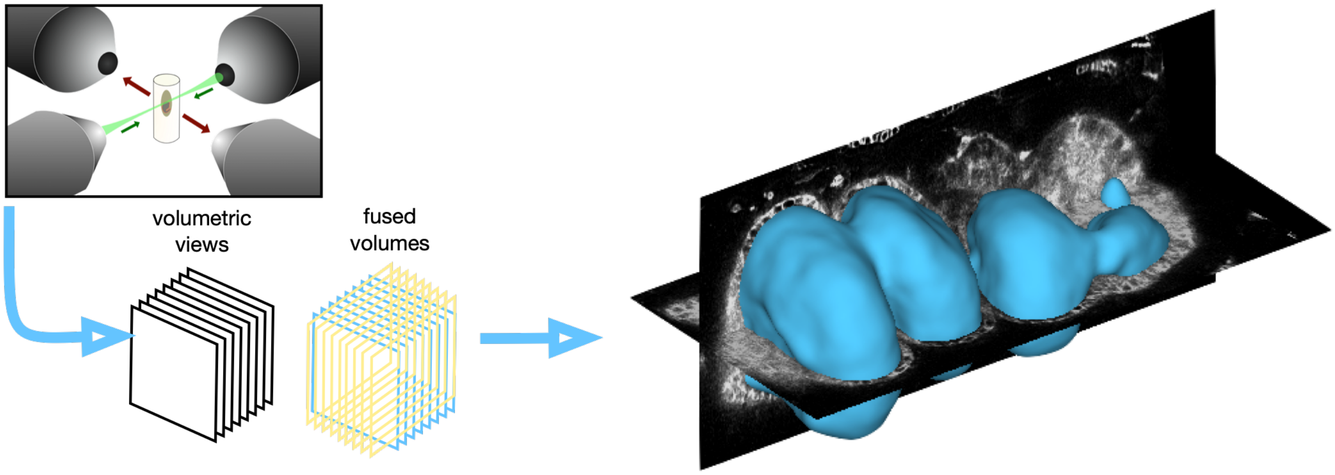


Supplementary Figure S1. **Multi-view light-sheet microscopy enables volumetric imaging.**

### III. MORPHOLOGICAL ALIGNMENT OF GUT SHAPE DYNAMICS

To morphologically align gut shape evolution across different embryos, we match the timepoint $t'$ of each sequence of shapes to the timepoint $\tau$ of a 'reference' dataset. To match timepoints, we optimally register the midgut surface of embryo $i$ at timepoint $t'$, $S_i(t')$, to all surfaces of the reference, $S_0(t)$ using an iterative closest point scheme on a subsampling of the vertices in each surface. Each midgut surface $S_i(t')$ is sampled into a pointcloud with an areal density of $\sim$0.01 $\mu\text{m}^{-2}$, and we compute the rigid transformation that registers the point cloud of surface $S_i(t')$

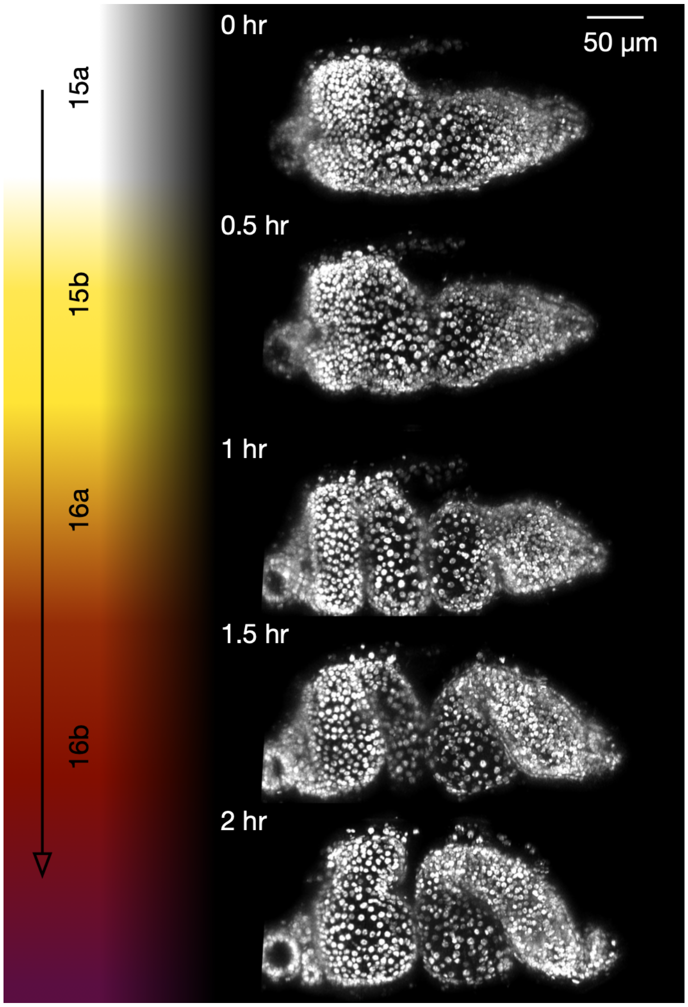


Supplementary Figure S2. **After deconvolution, 3D dynamic datasets offer subcellular resolution of the full organ.** Here, maximum intensity projections of left lateral views (left half of data volume) for visceral expression of *48Y GAL4* × *UAS-histone RFP* show organ shape change without cell proliferation. Some cells outside the gut driven by *48Y GAL4* are visible on the dorsal side.

with a point cloud of each midgut surface $S_0(t)$ in the reference dataset. Near the optimal time ($t = \tau$), the sum of squared residuals (distances) of paired points in the subsampled point clouds varies quadratically with reference time $t$. We identify the timepoint with the minimum sum of squared residuals as the morphological time $\tau$ of $S_i(t')$ and fit the nearby seven timepoints to a parabola to measure the uncertainty (colored bands in the main text Figure 1F. In addition to defining the morphological time for each timepoint of each embryo, this procedure also defines a rigid-body translation and rotation that optimally registers the surface $S_i(t')$ to $S_0(\tau)$. Example cross-sections of these morphologically aligned shapes for four timepoints ($\tau = 0, 0.5, 1, 1.5$ hr) are shown in Fig. S3.

### IV. PARAMETERIZATION OF THE ORGAN SHAPE

To quantify geometry of organ shape and deformation, we built an analysis package called TubULAR [8]. While details of the package are included in [8], we provide a high level overview of the initial steps of the pipeline here to discuss how the organ surface can be parameterized with a coordinate frame using 'intrinsic' AP-DV coordinates based on conformal mapping of the curved shape to a flat plane.

We begin by extracting the organ shape using Morphsnakes level set analysis [9, 10] on the output of an iLastik training [11] against membrane (or nuclei, actin, or myosin), such that the yolk within the midgut is predicted to fall into a single classification that is distinct from the midgut tissue. We aligned the organ into the AP-DV coordinate system by (1) training on the junction with the proventriculus on the anterior end in iLastik for a timepoint at the

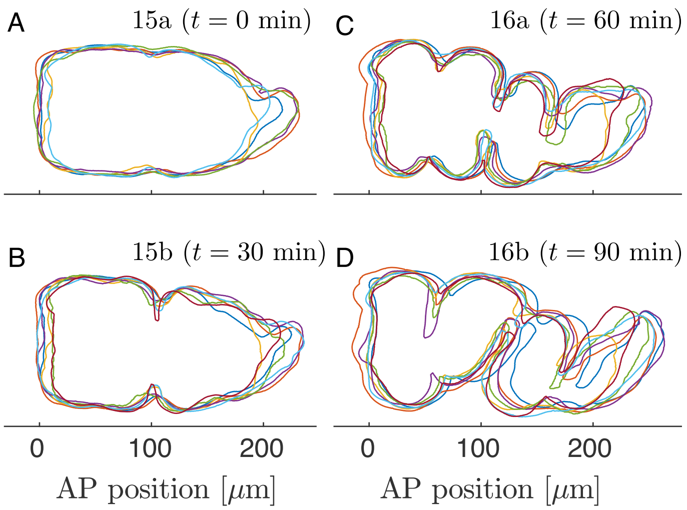


Supplementary Figure S3. **Computationally aligned midgut shapes demonstrate the degree of reproducibility during midgut morphogenesis.**

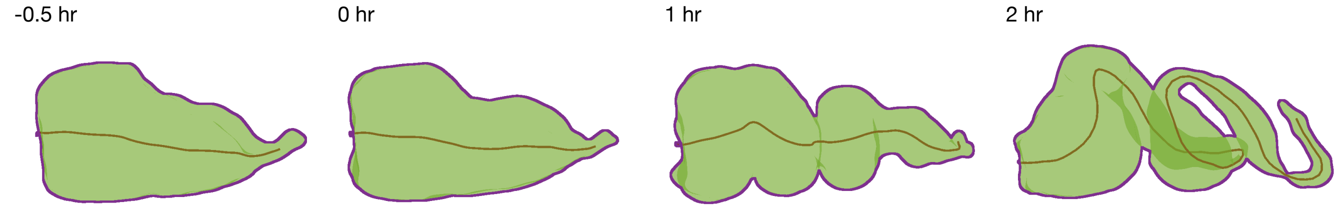


Supplementary Figure S4. **A centerline (brown) measures an effective length of the organ.** We compute the centerline (brown curve) using the TubULAR package for extracted shapes (green with purple boundary for clarity). The cumulative length of these curves defines the effective length of the organ reported in the main text.

onset of folding, (2) training on the posterior-most surface of the interface between midgut yolk and apical posterior midgut for the same timepoint, (3) training on the seam of midgut dorsal closure in iLastik for the same timepoint, (4) rotating the surface so that the anterior and posterior points align with the $x$ axis and the ray connecting the dorsal point to the nearest point along the anterior-posterior line segment aligns with the $-z$ direction. This ensures anterior to posterior corresponds to $x$, ventral to dorsal corresponds to $z$, and rightward lateral dimension corresponds to $y$. We then cut small anterior and posterior endcaps from this surface by (1) training on the junction with the proventriculus on the anterior end in iLastik for all timepoints, (2) training on the posterior-most surface of the interface between midgut yolk and apical posterior midgut for all timepoints, (3) computing the centers of mass of the iLastik output for anterior and posterior, (4) removing a topological disk enclosing any contiguous surface triangles within a set distance from the centers of mass. These cylindrical cut meshes are then reparameterized according to a harmonic map minimizing Dirichlet energy [8] as a cylindrical orbifold-Tutte embedding. After a geometric stabilization step to remove jitter by allowing longitudinal ($u$) displacements and circumferential ($v$) rotations of each curve $u$ =constant in the parameterization plane by optimally matching the (3D embedding) coordinates of the previous timepoint's discretized $uv$ coordinate grid, we smooth meshes in time with a triangular-pulse kernel of width $\pm 2$ minutes to define our $(s, \phi, t)$ coordinate parameterization (Fig. S5).

### V. MEASUREMENTS OF ORGAN CENTERLINE

We compute the centerline using the TubULAR package [8], wherein the organ is divided into circumferential 'hoops' based on its isothermal coordinates (Fig. S4). Hoops for which $s$ =constant define an effective circumference for increments along the length of the organ, and the average 3D position of each hoop defines its centerline point. Connecting mean points of adjacent hoops along the length of the organ defines the centerline of the object (brown curve) whose length is reported in the main text Figure 1G. Effective radii shown in Figure 1H of the main text are computed as distances from the circumferential hoops to the centerline.

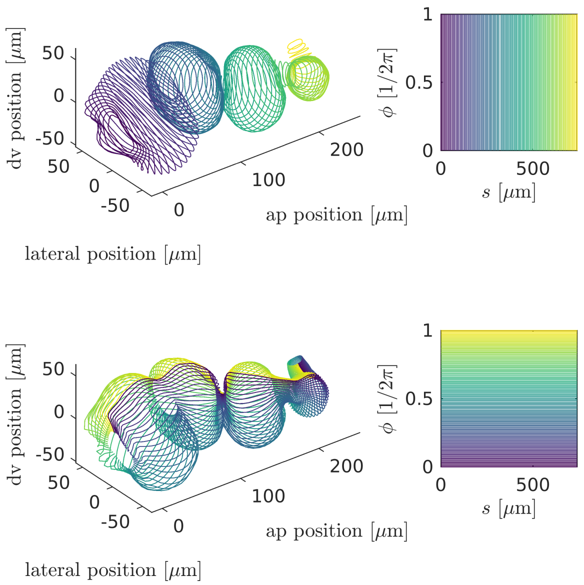


Supplementary Figure S5. **A surface-Lagrangian coordinate system parameterizes the gut surface by a 2D net $(s, \phi)$ to facilitate tissue velocity and deformation measurements.**

### VI. CIRCULARITY OF CONSTRICTIONS

We compute the transverse cross-sectional plane for each constriction for the main text Figure 1I. The constricting ring is obtained by first dividing the $(s, \phi, t)$ coordinate parameterization into curves of constant $s$, which are circumferential rings on the organ. We then define an effective radius as the average distance from each point in a uniform sampling of the ring to the centerline, which is composed of the mean positions of all circumferential rings.

Then constriction locations are identified as circumferential rings whose effective radii $r(s)$ are local minima (so that $\partial_s r(s) = 0$). Local minima in effective radius are tracked starting at the onset of folding forward in time to define constriction locations. Before the onset of folding, presumptive constriction locations are inferred by back-tracing the onset location to earlier timepoints.

### VII. ENDODERMAL CELL SEGMENTATION AND SHAPE CHANGE

Using a single slice of the gut surface projected into stabilized $(s, \phi)$ pullback coordinates, we segmented 600-1300 cells per timepoint (Fig. S6) using a semi-automated procedure:

1. We first perform adaptive histogram equalization over patches of the pullback containing several cells in width.

2. We then perform two passes of morphological image reconstruction (see MATLAB's `imreconstruct` function) punctuated by morphological dilation and erosion steps.

3. The result is binarized and skeletonized via a watershed algorithm.

4. We overlay this skeleton on the original image to enhance the membrane contrast, convolve with a narrow Gaussian (with a standard deviation of $\sim 0.2\%$ of organ length), and pass the result through the previous three steps.

This gives us an estimate for the image segmentation. We then manually correct any spurious segmentation artifacts in GIMP [12] by overlaying the segmentation with the original pullback images. To resolve some ambiguous cell junctions, we toggle between a maximum intensity projection of several microns along the surface normal direction and a single slice of the endodermal cell layer near the apical side (about 2.5$\mu$m beyond the apical side).

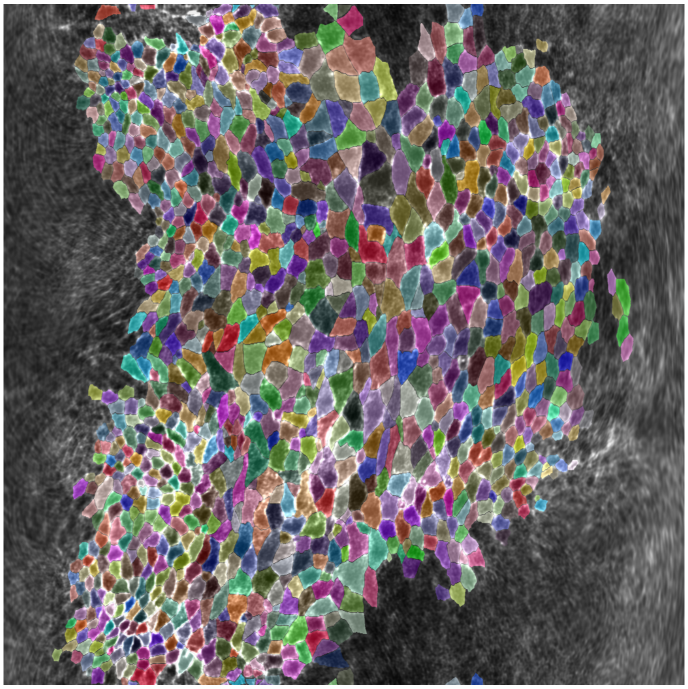

Supplementary Figure S6. **Example segmentation in the pullback plane at a time near the onset of folding.** Each cell polygon is given a random, distinguishable color to demonstrate the segmentation quality.

We compute cell anisotropy by finding segmented cell shapes in 2D, embedding those polygons in 3D, projecting each cell onto a local tangent plane, and measuring the moment of inertia tensor of this polygon in a suitable (material) coordinate system. This procedure is shown schematically in Fig. S7. We then embed the Lagrangian coordinate directions $(\hat{s}, \hat{\phi})$ from a conformal mapping of the whole organ at the onset of the initial (middle) constriction $t = t_0$ onto the cell's centroid in 3D (in the deformed configuration at time $t \neq t_0$). The moment of inertia tensor for the cell polygon is expressed in the local coordinate system from the embedded $(\hat{s}, \hat{\phi})$ directions in the local tangent plane of the tissue. The eigenvalues $I_1$ and $I_2$ of the moment of inertia tensor and their associated eigenvectors then provide an effective ellipse for the cell with orientation $\theta$ with respect to the local $\hat{\mathbf{s}}$ direction and an aspect ratio $\sqrt{I_1/I_2}$. Fig. S8 shows the raw data of these measurements without computing statistics.

We then wish to average the cellular anisotropy over the organ surface to report a mean, standard deviation, and standard error for both the cellular aspect ratio and cell orientation. In this averaging, we weight each cell's contribution to the observable (aspect ratio $a/b$ or orientation $\theta$) by its area, so that all material points on the organ are given equal weight. The results reported in Figure 2C and D in the main text are the weighted mean and weighted standard deviation for each distribution. The weighted mean of the aspect ratio $a/b$ orientation $\theta$ is therefore computed via

$$\langle a/b \rangle = \frac{\sum_{i=1}^{N} A_i a_i / b_i}{\sum_{i=1}^{N} A_i} \tag{1}$$

$$\langle \theta \rangle = \tan^{-1}\left[\frac{\sum_{i=1}^{N} A_i \sin\theta_i}{\sum_{i=1}^{N} A_i \cos\theta_i}\right], \tag{2}$$

where $A_i$ is the area of the $i^{\text{th}}$ cell of $N$ cells. We note that we obtain similar results by weighting each cell equally.

We obtain standard errors by bootstrapping. In detail, we subsample our collection of measurements, compute the weighted mean for the subsample, and repeat with replacement 1000 times. The variance of these 1000 means decreases in proportion to the number of samples $n$ included in our subsampling, so that $\sigma^2_{\overline{x}}(n) = \tilde{\sigma}^2_{\overline{x}}/n + \sigma^2_0$. Fitting for $\sigma^2_{\overline{x}}$ across 50 values of $n$ ($N/4 < n < N$) and evaluating this fit for $n = N$ gives an estimate for the standard error on the mean $\sigma_{\overline{x}} = \sqrt{\sigma^2_{\overline{x}}}$.

In Fig. S9, we show measurements of endodermal cell anisotropy for each of the four chambers separately. Though different chambers have varying degrees of initial anisotropy, the same motif of cell shape change is seen in each. Cells are initially oriented with their long axes along the circumferential direction. Over time, their aspect ratio approaches

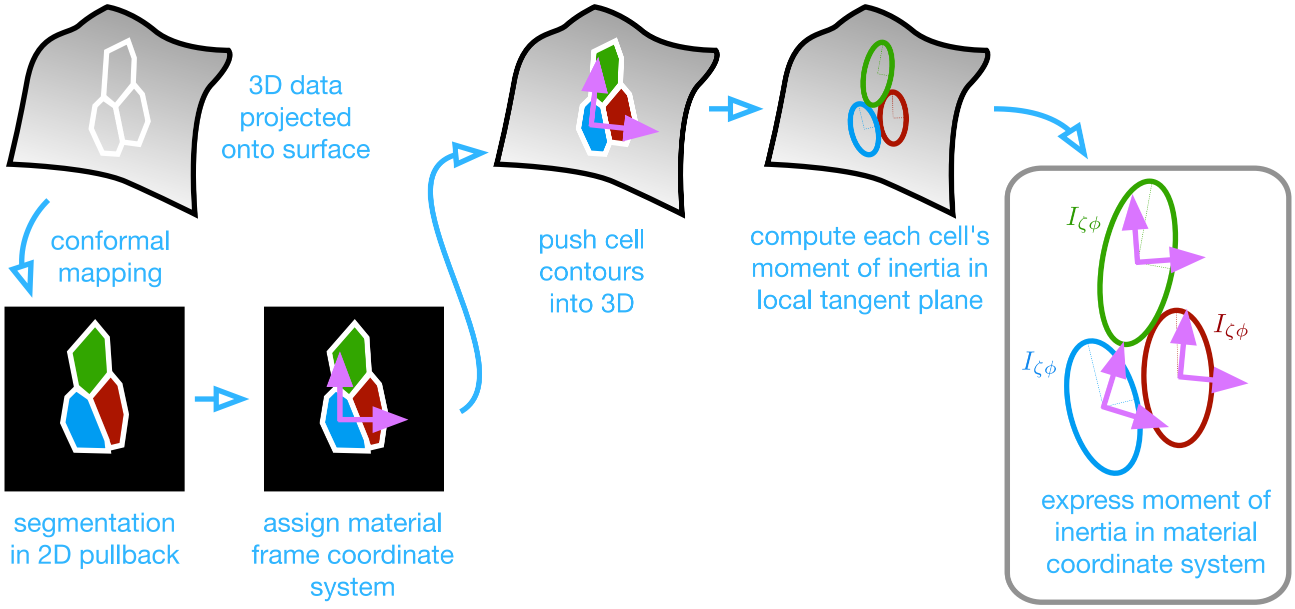


Supplementary Figure S7. **Our cell measurement pipeline utilizes conformal mapping to a 2D pullback plane for segmentation and re-embedding cells into local tangent planes for shape quantification.**

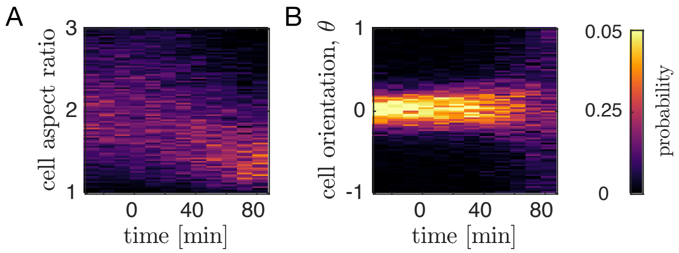


Supplementary Figure S8. **Raw histograms of cell shape changes over time.** *(A)* Cell aspect ratios decrease and condense close to 1 (isotropic shapes) by ∼80 minutes after the onset of the first (middle) constriction. *(B)* During this time, the cells do not rotate, as evidenced by the sustained peak in cell orientation near zero. At late times, when the cells are nearly isotropic, the orientation becomes less clearly defined and the distribution broadens.

1, in which case their orientation is not well defined, and the orientation either broadens or switches to 0 or $\pi$, meaning that the cells are then oriented with their long axes along directed along the organ's AP axis. In chamber 4, cells are notably less anisotropic at the onset of folding. The cell shapes in this chamber hit the isotropic limit $a/b \approx 1$, within cell-to-cell variation), and the orientation of the the long axis $a$ of the cells switches from $\pi/2$ to approximately zero, indicating alignment of the long axis with the organ's intrinsic anterior-posterior axis.

### VIII. SINGLE-CELL TRACKING

We tracked 175 cells from $-27$ min $< t <$ 83 minutes of development in a *48Y GAL4* × *UAS-CAAX::mCh* embryo imaged using muSPIM. First, we segmented the same 175 cells in the first chamber of the gut every two minutes using the same procedure as in the previous section. We tracked their positions over time using iLastik manual tracking workflow using 2D pullback projections. From these segmented polygons, we project back into 3D onto the gut surface and measure the cell areas in a local tangent plane for Figure 2E in the main text.

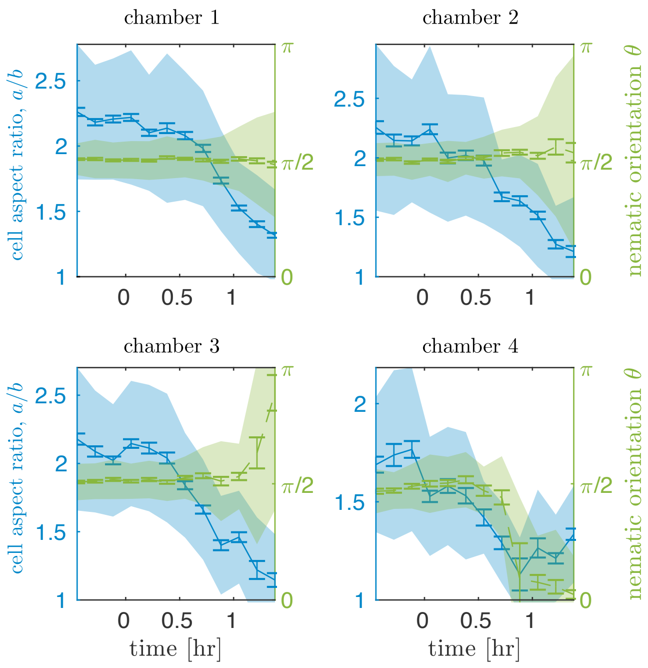


Supplementary Figure S9. **Endodermal cell shape change statistics show similar behavior across chambers.** Cell aspect ratios (blue) decrease during constrictions in all chambers. Cell orientations do not exhibit rotations, but do change toward zero or $\pi$ in the posterior chambers 3-4 once the cells become nearly isotropic ($a/b \approx 1$). Colored bands denote standard deviations weighted by cell area, and tick marks denote standard errors on the mean.

### IX. QUANTIFICATION OF TISSUE DEFORMATION

To compute a coarse-grained tissue velocity over the gut surface, we again used the TubULAR package [8]. This resource enables velocimetry and discrete exterior calculus measurements on the evolving surface. The result is a fully covariant measurement of the compressibility and shear of the tissue spanning the whole organ.

Briefly, given our $(s, \phi, t)$ coordinate system defined in the TubULAR pipeline, we then run particle image velocimetry (PIV) using PIVLab [8, 13, 14] and map tissue velocities in the domain of parameterization to the embedding space. Geometrically, displacement vectors $\mathbf{v}$ extend from one $\mathbf{x}(s_0, \phi_0, t_0)$ coordinate in 3D on the surface at time $t_0$ to a different $\mathbf{x}(s_1, \phi_1, t_1)$ coordinate on the deformed surface at time $t_1$. When $t_0$ and $t_1$ are adjacent timepoints, this define the 3D tissue velocity at $\mathbf{x}(s_0, \phi_0, t_0) = \mathbf{v}(s_0, \phi_0, t_0)/(t_1 - t_0)$. We decompose the velocity into a tangential component $\mathbf{v}_t$ and a normal component $\mathbf{v}_n = v_n\hat{\mathbf{n}}$ for measuring divergence via discrete exterior calculus [8] and for measuring bending $2Hv_n$, where $H$ is the mean curvature obtained via computing the Laplacian of the mesh vertices in (embedding) space: $\Delta\mathbf{X} = 2H\hat{\mathbf{n}}$.

We make sense of divergence and bending and interpret their difference as the local area growth rate by the following argument. The surface changes according to its tissue velocity, which has tangential and normal components $\mathbf{v} = \partial_t\mathbf{X} = v^i\mathbf{e}_i + v_n\hat{\mathbf{n}}$. The shape of the surface is encoded by the metric $g_{ij} = \partial_i\mathbf{X}\cdot\partial_j\mathbf{X}$, which describes lengths and angles measured in the tissue, and by the second fundamental form $b_{ij} = \partial_i\partial_j\mathbf{X}\cdot\hat{\mathbf{n}} = -\partial_i\mathbf{X}\cdot\partial_j\hat{\mathbf{n}}$, which contains information relating to both intrinsic and extrinsic measures of surface curvature. The time rate of change of the metric is determined by the superposition of velocity gradients and normal motion where the surface is curved:

$$\dot{g}_{ij} = \nabla_i v_j + \nabla_j v_i - 2v_n b_{ij}. \tag{3}$$

Here, $\nabla$ denotes the covariant derivative operator defined with respect to the embedding metric $\mathbf{g}$. The covariant mass continuity equation gives [15]

$$0 = \frac{D\rho}{Dt} + \frac{\rho}{2}\mathrm{Tr}[\mathbf{g}^{-1}\dot{\mathbf{g}}] \tag{4}$$

$$= \frac{D\rho}{Dt} + \rho\nabla\cdot\mathbf{v} - \rho 2v_n H, \tag{5}$$

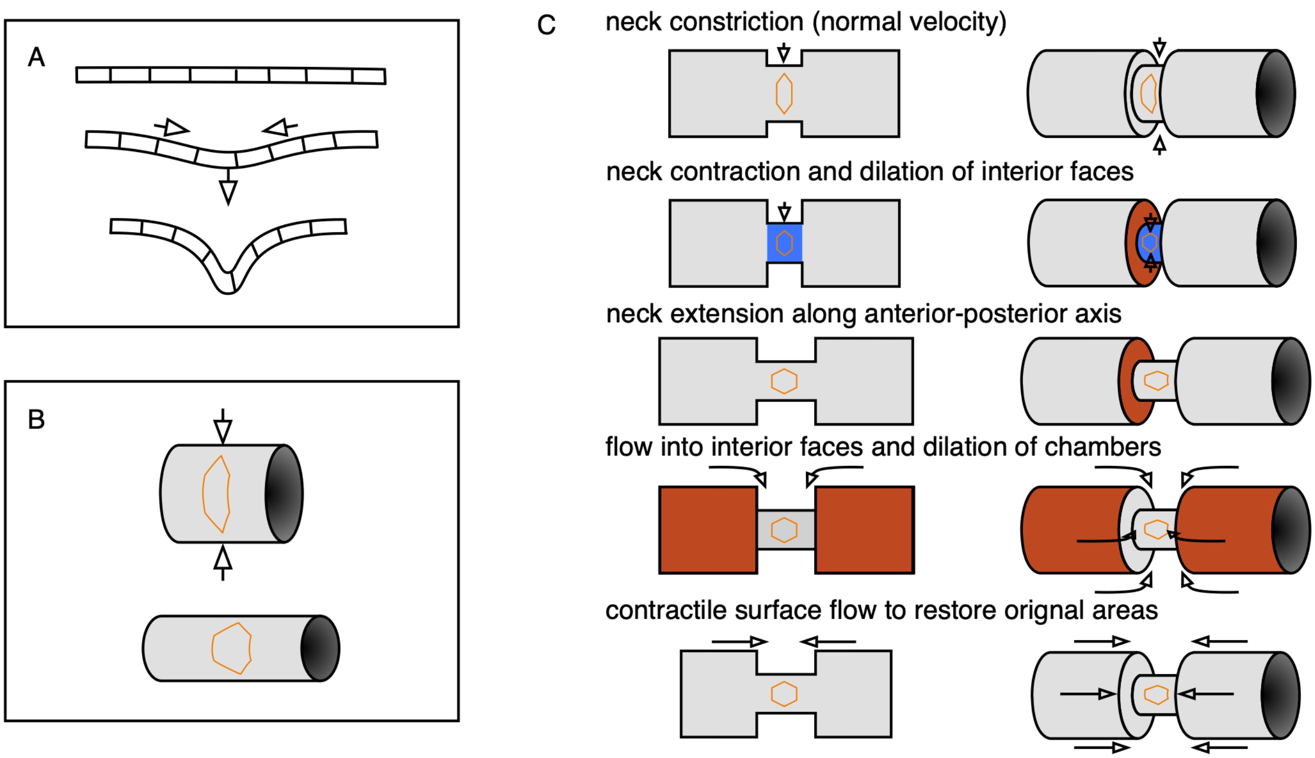


Supplementary Figure S10. **Three-component decomposition of kinematics elucidates coupling between compressibility and convergent extension.** *(A)* An initially flat incompressible sheet demonstrates kinematic coupling between contraction and bending to preserve cells' 2D areas. *(B)* Constriction couples to convergent extension via curvature of the surface. As a tube constricts, the tube elongates in order to preserve surface area. As a result, cells change their aspect ratio and undergo convergent extension. *(C)* The two effects couple in a pinching tube. The effects can be decomposed in the case of a tube with a step-like constriction composed of deformable cells whose areas shall not change. Constriction of the narrow tube via inward normal velocities would decrease the neck area (blue), so the neck extends to keep its area fixed. This is convergent extension. The faces are now dilated, triggering flow into the interior faces and contractile surface flows to raise the lower the areas of the dilated tissue. In this way, convergent extension is linked to incompressibility, which couples bending and contractile flows.

where $\rho$ is the mass density in the physical embedding, and the material derivative is $D\rho/Dt = \partial_t \rho + \rho(\nabla \cdot \mathbf{v}_\parallel) + \mathbf{v} \cdot \nabla \rho$. Incompressibility ($D\rho/Dt = 0$) then implies

$$\nabla \cdot \mathbf{v} = 2Hv_n. \tag{6}$$

### X. MINIMAL INGREDIENTS DEMONSTRATE GEOMETRIC INTERPLAY BETWEEN COMPRESSIBILITY AND SHEAR

A nearly incompressible sheet, such as a sheet of paper, naturally demonstrates the kinematic coupling between contraction ($\nabla \cdot v_t < 0$) and bending ($2Hv_n$). Contracting such a sheet as in Fig. S10A leads to out of plane bending to preserve surface area of the sheet. This out-of-plane deformation leaves cells unchanged in their aspect ratio: no in-plane deformation is necessary.

If the sheet is curved into a tube (so that mean curvature is nonzero, $|H| > 0$), then constricting an sheet (with inward velocity $v_n > 0$) can generate deformation in the local tangent plane of the sheet. Such incompressibility couples to initial curvature to generate shear deformation. For example, an incompressible sheet of paper glued into a cylinder along one of its edges cannot be pinched in this fashion without crumpling or folding. An elastic sheet, however, can be deformed in this manner even if local areas of material patches are required not to change. In particular, the sheet may stretch along the long axis while constricted circumferentially, such that a circular material patch is transformed into an elliptical patch with the same area, as in Fig. S10B.

Finally, these two effects are coupled in the case of the pinched cylinder. In a given snapshot with an existing localized constriction, we can schematically understand the three ingredients by considering the pinched cylinder

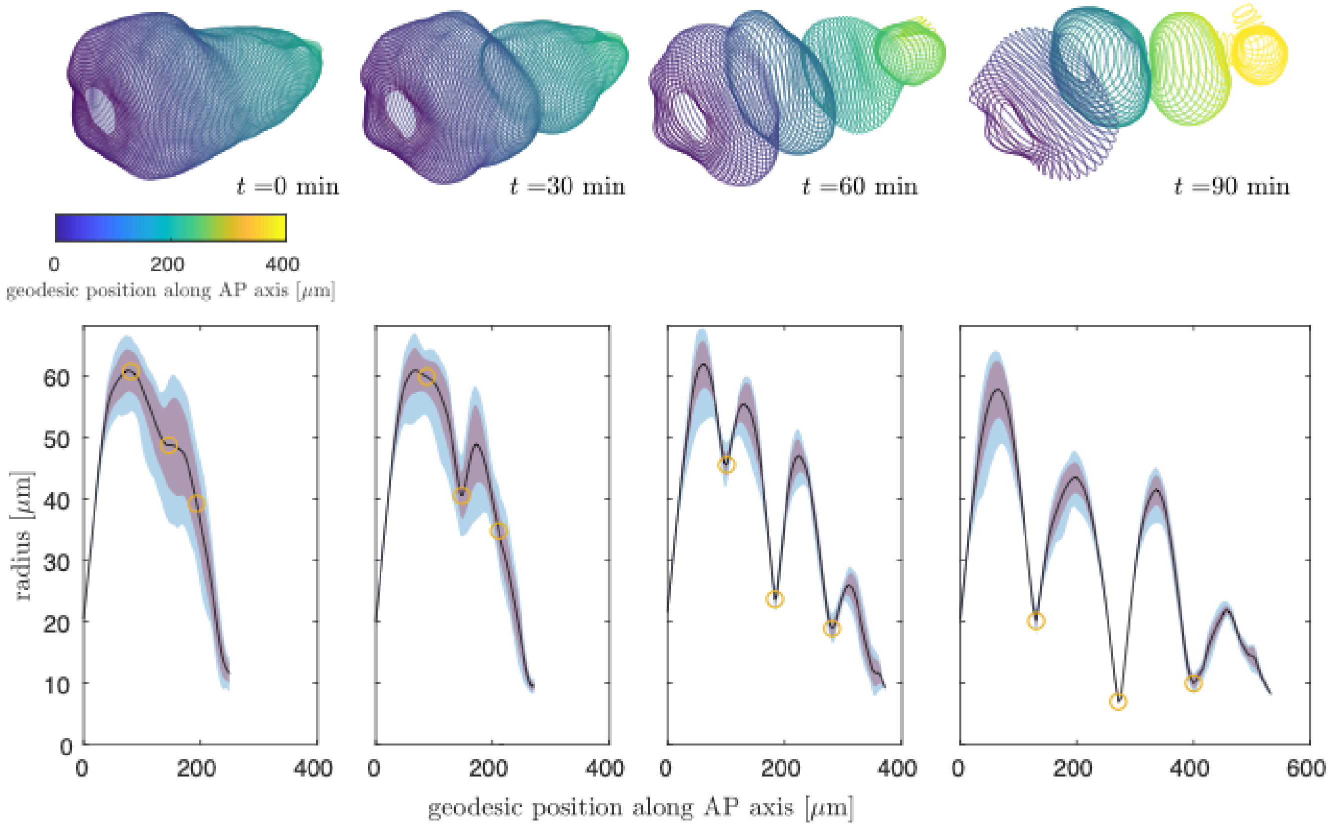


Supplementary Figure S11. **Cross-sectional radii tighten near constrictions, reflecting the circular shape of constriction cross-sections for a representative embryo.** Via whole-organ surface parameterization, we compute a cross-sectional width of the organ (effective radius) as a function of length from the anterior end to anterior-posterior position along the organ surface ($s$), which we compute as the average length along the surface from the anterior to uniformly-sampled points in a ring of constant $\phi$ in the conformal pullback space $(s, \phi)$. Intuitively, this is the average path length required to travel on the surface from the anterior face to a given location while tracing out a curve of constant $\phi$, which are geodesics at the onset of the first constriction. That is, $s = \langle \int_{\text{anterior}}^{\zeta} ds' \rangle$, where the average is taken along the circumferential 'hoop' of constant $\zeta$. The purple band represents the standard deviation of effective radii along each circumferential hoop ($\zeta$ =constant), while the light blue band reflects the spread between the maximal and minimal values of effective radius at each hoop. Yellow circles mark the minima of the effective radii at each constriction. The narrow distribution in distances from the centerline to different surface points at each constriction reflects the circular cross-sectional shape during constriction.

with a step-wise indentation shown in Fig. S10C. First, active stresses in the neck constrict the neck, decreasing the surface area of the neck (blue) and dilating the interior faces (red). In order to restore the surface area of the cells in the neck, the length may increase, resulting in extension along the long axis of the tube. In tandem, to combat the dilation in the interior faces, cells flow into the constriction from the chambers. Note that if all three steps are instantaneously coupled, the order could be reversed to accomplish the same outcome without change in observable behavior: contractile surface flows could increase the density of cells in the interior faces, which leads to neck constriction to restore cell density in the faces and results in convergent extension of the neck.

We note also that when the constriction is broad and/or shallow, such that the mean curvature is positive everywhere (cylinder-like, $H > 0$), then inward motion of the incompressible tube causes an extensile surface flow ($\nabla \cdot \mathbf{v}_{\parallel} > 0$), rather than a contractile one. This is a qualitative difference between broad or uniform constrictions of a tube and localized constrictions such as those seen in the midgut. In principle, we predict a crossover between the two modes of behavior during the very onset of constriction in our system – from positive to negative divergence as the curvature changes sign. This is a subtle and transient feature, if present at all, given the large radius of the midgut compared to the axial length of the constrictions and non-uniform initial curvature of the midgut before constrictions begin.

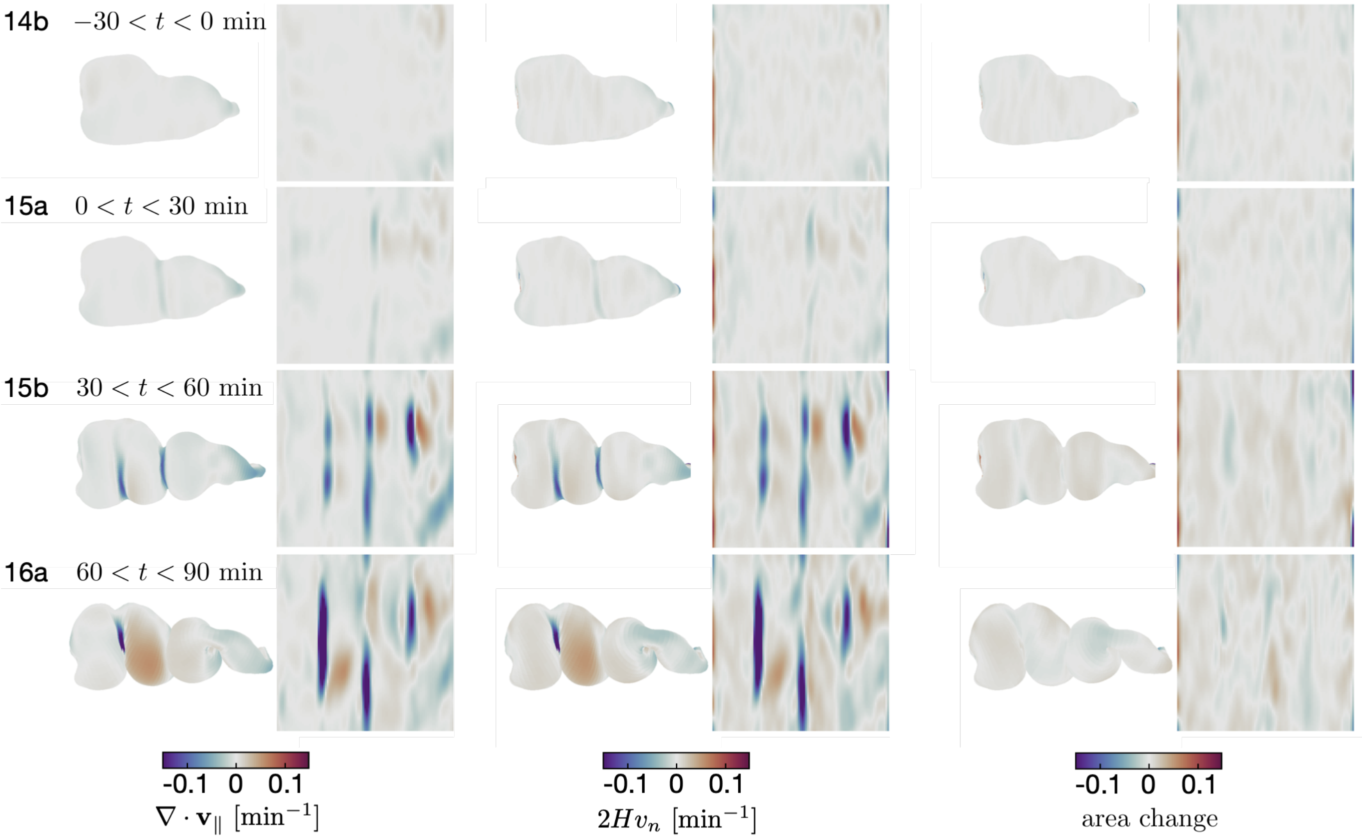


Supplementary Figure S12. **Contraction (and dilation) patterns are tightly coupled to bending throughout midgut constrictions, indicating a nearly incompressible tissue behavior.** Resulting divergence measurements from discrete exterior calculus and bending measurements on an representative embryo, plotted both on the midgut surface in 3D in pullback coordinates, show strong correlation. The difference of the two patterns gives the local area change (right columns). Each image is the average of patterns over a 30 minute timespan.

### XI. TISSUE SHEAR

We employ a geometric method of tissue-scale shear quantification that accounts for both the shear due to the changing shape of the gut and the shear due to the material flow of cells along the dynamic surface. The first step in the pipeline is to establish consistent material coordinates for all times, i.e. labels for parcels of tissue that follow those parcels as they move and deform. We prescribe these labels at the onset of folding by endowing a cylindrical 'cut mesh' of the organ's surface at the that time with a planar parameterization. The cut mesh is first conformally mapped into a planar annular domain, $\{||\vec{\mathbf{x}}|| : r \leq ||\vec{\mathbf{x}}|| \leq 1\}$, using a custom Ricci flow code included in our TubULAR package [8]. Fixing the outer radius of the annulus to 1, the inner radius $r$ is a conformal invariant that is automatically determined from the geometry of the organ. Taking the logarithm of these intermediate coordinates then defines a nearly rectangular domain, with a branch cut identifying the top and bottom horizontal edges of the domain in such a way the the cylindrical topology of the cut mesh in 3D is fully respected. The coordinates in this domain are taken to be the 'intrinsic' material (or Lagrangian) coordinate system, $(\tilde{s}, \tilde{\phi})$. This conformal parameterization is, by construction, totally isotropic and is therefore a suitable reference against which to measure all subsequent accumulation of anisotropy in the tissue.

In order to recapitulate the material flow of the tissue, these coordinates are advected in the plane along the flow fields extracted using PIV [13, 14] and then mapped into 3D at each time point. This mapping defines a deformed mesh whose induced metric tensor, $\mathbf{g}' \equiv \mathbf{g}(t)$, can be computed relative to the material coordinates. All anisotropy in the mapping is encoded by the complex Beltrami coefficient, $\mu(t)$, defined in terms of the components of the time-dependent metric tensor

$$\mu(t) = \frac{g'_{11} - g'_{22} + 2i\, g'_{12}}{g'_{11} + g'_{22} + 2\sqrt{g'_{11}\, g'_{22} - {g'_{12}}^2}}. \tag{7}$$

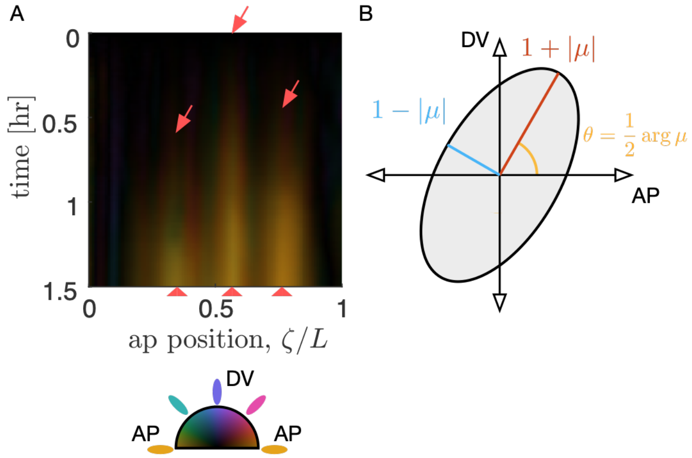


Supplementary Figure S13. **Tissue shear, generated by 3D convergent extension during constrictions, is captured in the Beltrami coefficient – a local measure of anisotropic, area-preserving deformation.** *(A)* Kymograph of a representative embryo's tissue deformation shows the emergence of oriented shear near folds. Here, the organ is parameterized by the anterior-posterior (AP) position, $\zeta$, at the onset of folding ($t = 0$). Folds begin to appear at times and locations marked by red arrows and continue to deepen. A measure of tissue deformation, known as the Beltami coefficient, is averaged along the circumferential (DV) direction and plotted at the anterior-posterior position in the material frame, so that the deformation of advected tissue patches are compared to their original shape at $t = 0$. Color denotes the orientation of the anisotropic shear deformation in the material frame, which is a conformal map of the (cut) organ tube onto a rectangle. Shear which extends along the intrinsic AP axis is denoted by orange color, and this color dominates the kymograph: the deformation is globally aligned to extend along the longitudinal AP axis (and contract along the circumferential axis). *(B)* The Beltrami coefficient $\mu$ is defined as the amount of area-preserving shear transforming a circle into an ellipse with aspect ratio $1 + |\mu|/(1 - |\mu|)$ at an angle $\arg\mu/2$, which is denoted by color in the kymograph.

As illustrated in Fig. S13, $\mu$ describes how an initially circular infinitesimal patch of tissue is deformed into an elliptical patch under the action of the material mapping. The argument of $\mu$ described the orientation of this ellipse. The magnitude $|\mu|$ is related to the ratio, $K$ of the lengths of the major axis of this ellipse to its minor axis by

$$K = \frac{1 + |\mu|}{1 - |\mu|}. \tag{8}$$

When $|\mu| = 0$, the material mapping is isotropic, i.e. a circular patch of tissue remains circular under the mapping. Note that $|\mu| < 1$ and therefore provides a bounded description of both the magnitude and orientation of material anisotropy in the growing surface.

### XII. RELATIVE MOTION BETWEEN LAYERS

To characterize relative motion between layers, we tracked 375 endodermal and 81 muscle nuclei in the same *w,Hand>GAL4;UAS-Hand:GFP;hist:GFP* embryo. Fig. S14 shows individual track measurements of relative displacement of initially-close nuclei pairs ($< 5$ $\mu$m apart at the onset of the first constriction). Two example tracks are highlighted in yellow and green. Fig. S15 shows additional statistics of the relative motion over time.

### XIII. OPTOGENETIC EXPERIMENTS

The *OCRL* and *UAS-RhoGEF2* constructs have been previously characterized [16, 17]. We provide schematics of their cellular mechanisms in Fig. S16. For optogenetic confocal microscopy experiments, we activated the optogenetic construct with continuous oblique illumination of a 470 nm LED at 6.2±0.1 mW/cm$^2$ power, in addition to periodic illumination with the 488 nm laser used to image the sample. Wild-type embryos developed normally under this illumination ($N = 18/19$). Variations by a factor of two in either the LED power or in the 488 nm laser power used to image the *GFP* channel did not result in differences in phenotype. For light sheet imaging, we illuminated with 488 nm laser line at 1 mW for 30 seconds once per minute.

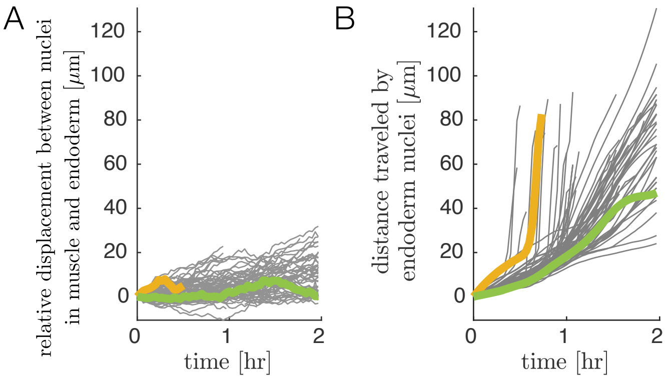


Supplementary Figure S14. **Relative motion of endodermal and mesodermal nuclei is small compared to motion of the tissue.** *(A)* By tracking 81 muscle nuclei and the nearest counterpart nucleus in the endoderm for each reveals only a gradual increase in geodesic distance (distance along the gut surface) over time. The mean displacement grows by $sim5$ $\mu$m per hour during folding on average, regardless of whether nuclei are located in deep folds (yellow curve) or on the surface of the gut chambers (green curve). *(B)* In contrast, the tissue deformation leads to large displacements of cells. We measure distances in embedding space along pathline trajectories suitably smoothed to remove contributions from noise and transient motions. Distance traveled for example tracks invaginating into deep folds (yellow curve) or translating on gut chambers (green curve) are highlighted to match panel (A).

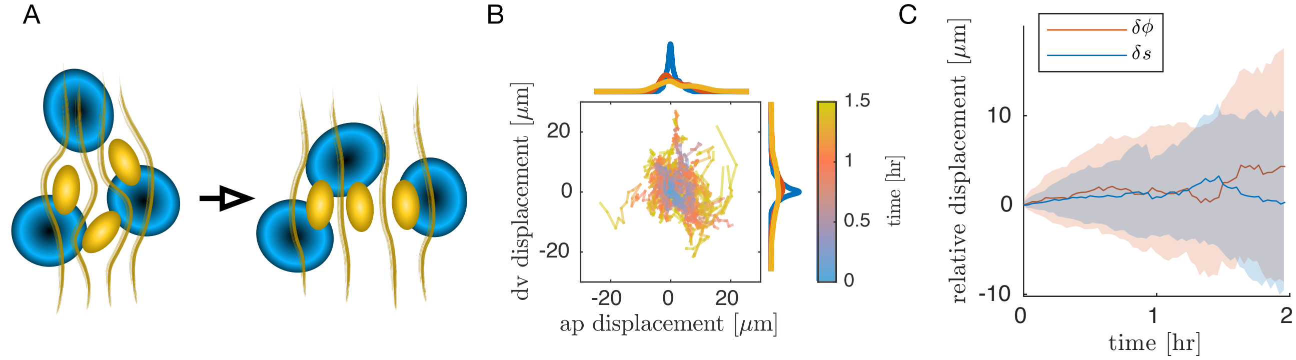


Supplementary Figure S15. **Muscle nuclei motion with respect to the endoderm is not coherent.** *(A)* Schematic of muscle nuclei configuration at early and late times. Muscle cells are positioned in bands along the AP axis and are initially clustered, such that each band is several cells wide along the circumferential axis. As constrictions form, circumferential muscle nuclei arrange in a nearly single-file configuration. *(B)* Motion of nuclei cells relative to the endodermal layer does not show strongly coherent (directional) motion, as demonstrated by individual tracks of relative displacement colored by timestamp. As before, the positions of 81 nuclei cells are measured relative to the center of mass of the endodermal nucleus that was nearest at the onset of folding. Distances are measured as geodesic lengths along the surface. Distances along the normal direction (through the thickness of the tissue) are ignored. That is, nuclear positions are projected along the thickness of the tissue onto the surface in which the endodermal nuclei reside. Histograms of accumulated displacements in either direction show the average over the first 30 minutes (blue), 30-60 minutes (red), and 60-90 minutes (yellow). *(C)* The same data shown in (B) is plotted as a distribution, with each component separated. The standard deviation of displacement coordinates (colored bands), either in the $s$ direction (along the folding AP axis in the material frame, blue) or the $\phi$ direction (along the circumferential axis in the material frame, orange) show an increase of $\sim 5$ $\mu$m per hour, with nearly zero mean displacement in either axis.

We quantified the endoderm cell shapes using a similar procedure as before. After deconvolution (Huygens Essential software), we perform 3D segmentation via a morphological snakes level set method on an iLastik pixel classification to carve out an approximate midsurface of the endoderm. We measured the endoderm shape dynamics for two-color *Antp-GAL4* $\times$ *UAS-OCRL* embryos held under continuous optogenetic activation from oblique illumination of a 470 nm LED at 6.2$\pm$0.1 mW/cm$^2$ power as before. For comparison, we additionally measured endoderm shapes in *Antp* mutant embryos with a membrane marker driven in the midgut endoderm ($w;48Y\ GAL4;Antp^{NS+RC3}$ $\times$ *UAS-CAAX::mCh*).

Fig. S17A shows representative snapshots of this segmentation procedure for a two-color *Antp-GAL4* $\times$ *UAS-OCRL* embryo at the onset of the first constriction and 40 minutes after the middle constriction began. Circumferential muscle localized near the anterior fold expresses the optogenetic construct (cyan band), while the endoderm is imaged using a ubiquitous membrane marker (grayscale). Image regions masked in semi-transparent gray are the deepest confocal plane acquired, while the rest of the image is a lateral view of the projected data on the segmented organ surface. Segmented endodermal cell polygons are colored by their aspect ratios. Cells are segmented in 2D and then projected into 3D for measurement of their aspect ratios. As shown in Fig. S17C, there is no significant difference between cell orientations in wild-type (blue), optogenetic mutants (red), and *Antp* mutants (yellow).

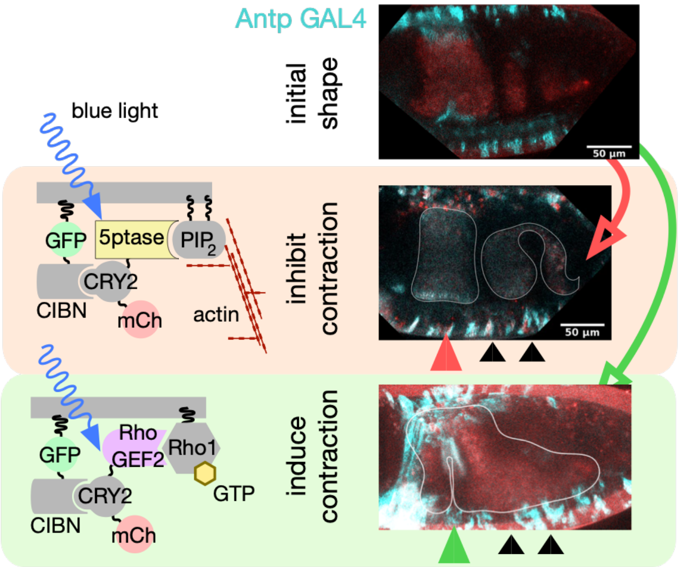


Supplementary Figure S16. **Confocal imaging of optogenetic mutants supplements the light sheet data shown in the main text.** Example sagittal section of a wild-type *Antp-GAL4* × *UAS-CIBN::GFP* (top) shows the anterior constriction forming in the middle of the *Antp* muscle band. Optogenetic inhibition of contraction in the *Antp* band abolishes the anterior fold, shown in sagittal section of a confocal stack (middle row). Optogenetically stimulating contraction in the *Antp*-expressing muscle band induces a constriction the anterior fold long before the constriction would normally appear, shown in sagittal section of a confocal stack (bottom row).

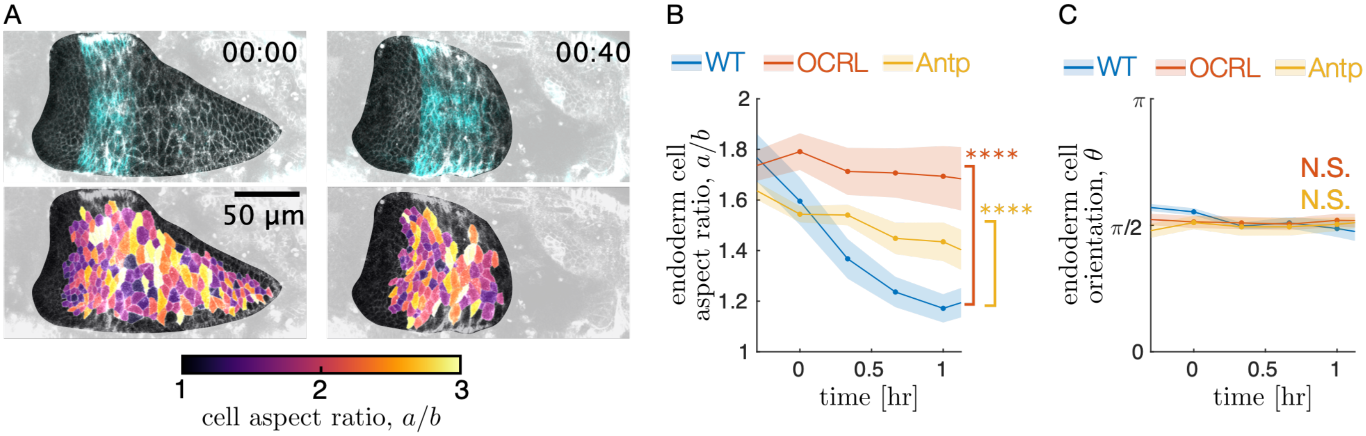


Supplementary Figure S17. **Optogenetic knockdown of muscle contractility inhibits endodermal shape changes, mimicking mutant behavior.** *(A)* Snapshots of single-cell shape measurements of embryos under optogenetic perturbations demonstrate that muscle contraction induces endoderm cell shape change. During optogenetic inhibition of muscle contractility in the *Antp* domain using *Antp-GAL4* × *w; UAS*>*CIBN::pmGFP; UAS-OCRL*, cell shapes in the interior two chambers (which remain as a single chamber in the optogenetic mutant) remain steady. *(B)* Endodermal cells undergo less shape change in both *OCRL* and *Antp* mutants, as reported in the main text, and as before, $**** = p < 0.0001$. *(C)* The endodermal cell orientation does not change significantly between conditions.

### XIV. WILD-TYPE CALCIUM DYNAMICS

We quantified calcium dynamics using the confocal microscopy (Leica SP8) of the live reporter *UAS GCaMP6s* driven by either the driver *Mef2-GAL4*, which is expressed across all muscles in the embryo, or *48Y GAL4*, which is expressed in the embryonic midgut both in endoderm and visceral muscles. We found that the two drivers yielded similar quantitative results for the anterior fold. Quantification of the posterior constriction was difficult to perform with *Mef2-GAL4* due to the presence of many fluorescent, non-midgut muscles near the changing focal plane location of the midgut constriction.

To measure transient calcium activity without bias from variations in ambient fluorescent intensity due to spatially-dependent scattering, we imaged three confocal stacks with 2.5-3 $\mu$m step size in rapid succession (9-10 seconds apart) and subtracted subsequent image stacks from each other according to

$$\delta I \equiv |I_1 - I_2| + |I_2 - I_3| + |I_1 - I_3|, \tag{9}$$

where $I_i = I_i(x, y)$ is the maximum intensity projection (projected across $dz \approx 30\mu$m) of the $i^{\text{th}}$ stack. Over such short timescales, motion of the midgut is small, but transient flashes of *GCaMP6s* are unlikely to span more than one acquisition. We then extract coherent features from $\delta I$ using a Gaussian blur followed by a tophat filter, and sum the resulting signal along the circumferential direction.

While we interrogated *GCaMP6s* activity using many views of the gut, the quantification was done using three standardized views. For the anterior constriction, we used a dorsal view, since out-of-plane effects (which could generate caustics as the fold deepens) are smallest on the dorsal side and since the midgut is nearest to the surface on the dorsal side. For the middle constriction, we used ventrolateral views since there is a line of sight absent of other muscles driven by *Mef2-GAL4* from this view. For the posterior constriction, we used a left lateral view for quantification.

To time-align the *GCaMP6s* experiments of the anterior and middle constrictions, we defined $t = 0$ as the first timestamp in which the constriction under observation showed localized bending along the AP axis. For characterization of the posterior constriction, the onset of folding was defined as the onset of fast phase of constriction, corresponding to when the ventral side of the gut visceral muscle had moved dorsally by $\sim$10 $\mu$m.

Fig. S18 shows kymographs of *GCaMP6s* dynamics averaged across biological repeats. In these kymographs, activity begins near the time when constrictions begin for each fold. In contrast, Fig. S19 shows delayed and suppressed *GCaMP6s* activity in *Antp* mutants compared to the wild-type behavior of sibling embryos that are not homozygous mutants for *Antp*.

### XV. CALCIUM ACTIVITY IN *ANTP* MUTANTS

To compare calcium activity in *Antp* mutants against wild-type dynamics, we computed $p$ values using a $z$-score measuring the difference between two populations: *Antp* heterozygotes (controls) and *Antp* homozygotes (mutants) as

$$Z = \frac{\overline{\delta I}_{\text{mutant}} - \overline{\delta I}_{\text{control}}}{\sqrt{s^2_{\text{control}}/n_{\text{control}} + s^2_{\text{mutant}}/n_{\text{mutant}}}}, \tag{10}$$

where $\overline{\delta I}$ is the sample mean, $s_{\text{control}}$ and $s_{\text{mutant}}$ are the sample standard deviations, and $n_{\text{control}}$ and $n_{\text{mutant}}$ are the sample size. This score gives a single-sided $p$ value via

$$p = \frac{1}{2}\text{erfc}\left(-Z/\sqrt{2}\right), \tag{11}$$

where erfc is the complementary error function.

To quantify the difference in overall activity between mutants and heterozygotes, we first estimate the expected fluorescent intensity for a given embryo under the null hypothesis that all embryos, whether mutant or not, will have similar *GCaMP6s* activity. Since embryos vary in opacity, we normalized each heterozygous embryo according to a value dependent on its background fluorescent intensity measured in regions within the embryo but far (45-50 $\mu$m) from the site of the putative fold. The observed maximum fluorescent activity $\delta I$ correlated with this background signal with a correlation coefficient of 78% and a mean signal-to-background ratio of 5.1$\pm$0.5. We then normalized each embryo's time-averaged $\delta I = \delta I(x)$ as

$$\delta I \to \frac{\delta I - \delta I_{\text{bg}}}{\delta I_{\text{max}} - \delta I_{\text{bg}}}. \tag{12}$$

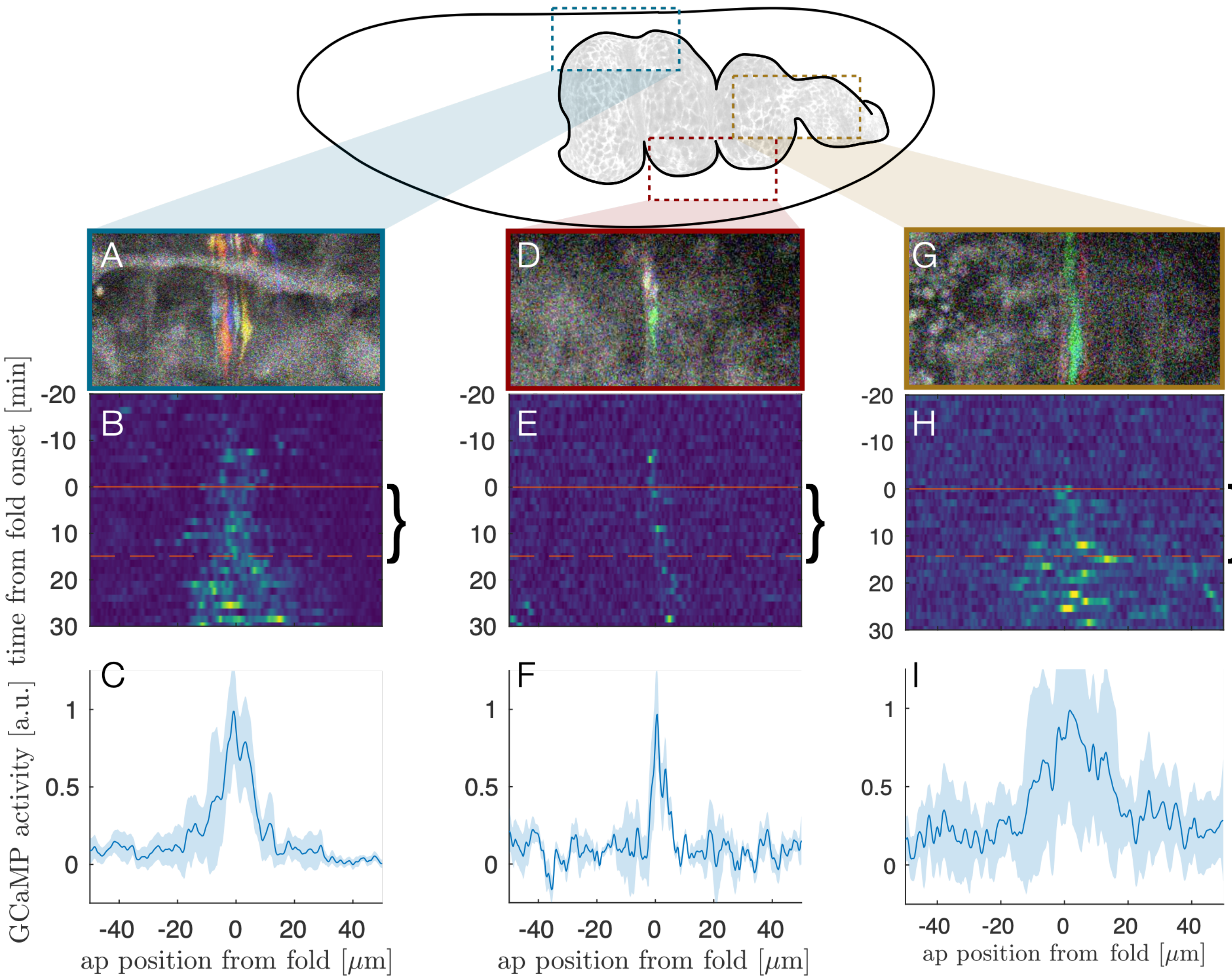


Supplementary Figure S18. **Kymographs of *GCaMP6s* dynamics show that calcium activity is initially localized in space to the site of folding and begins near the time of folding.** For each constriction, the transient signal is computed (colored signals in snapshots A, D, and G) and averaged across the DV direction into a space-time heatmap (B, E, and H). A time of $t = 0$ min for each panel corresponds to the time when localized constriction is visible in the bright-field channel at that constriction location and carries an uncertainty of $\pm 5$ minutes. The middle constriction (the sharpest fold, D-F) has the sharpest activity profile, and the posterior constriction (the posterior fold, G-I) has the broadest activity profile. The average activity during the first 15 minutes post folding onset is collapsed into the curves in (C), (F), and (I).

This enabled us to reduce the confounding influence of variation in optical density between embryos in the mutant analysis and compare absolute curves $\delta I$ rather than only their variation along the anterior-posterior axis.

### XVI. RNAI AND SERCA MUTANT ANALYSIS

To drive expression of RNAi or a dominant negative allele of *SERCA*, we administered heat shock by abruptly raising the temperature to 37°C using a stage-top incuabator (Okolab) and observing embryos staged such that they had not yet completed gut closure. The standard errors in the probabilities of successful folding are given by

$$SE = \sqrt{\frac{\hat{p}\,(1-\hat{p})}{N}}, \tag{13}$$

where $\hat{p}$ is the observed frequency of forming all three constrictions and $N$ is the number of samples of a given genotype (for ex, *Mef2-GAL4* $\times$ *UAS-SERCA.R751Q::mtomato*) measured in the experimental heat shock conditions. We note

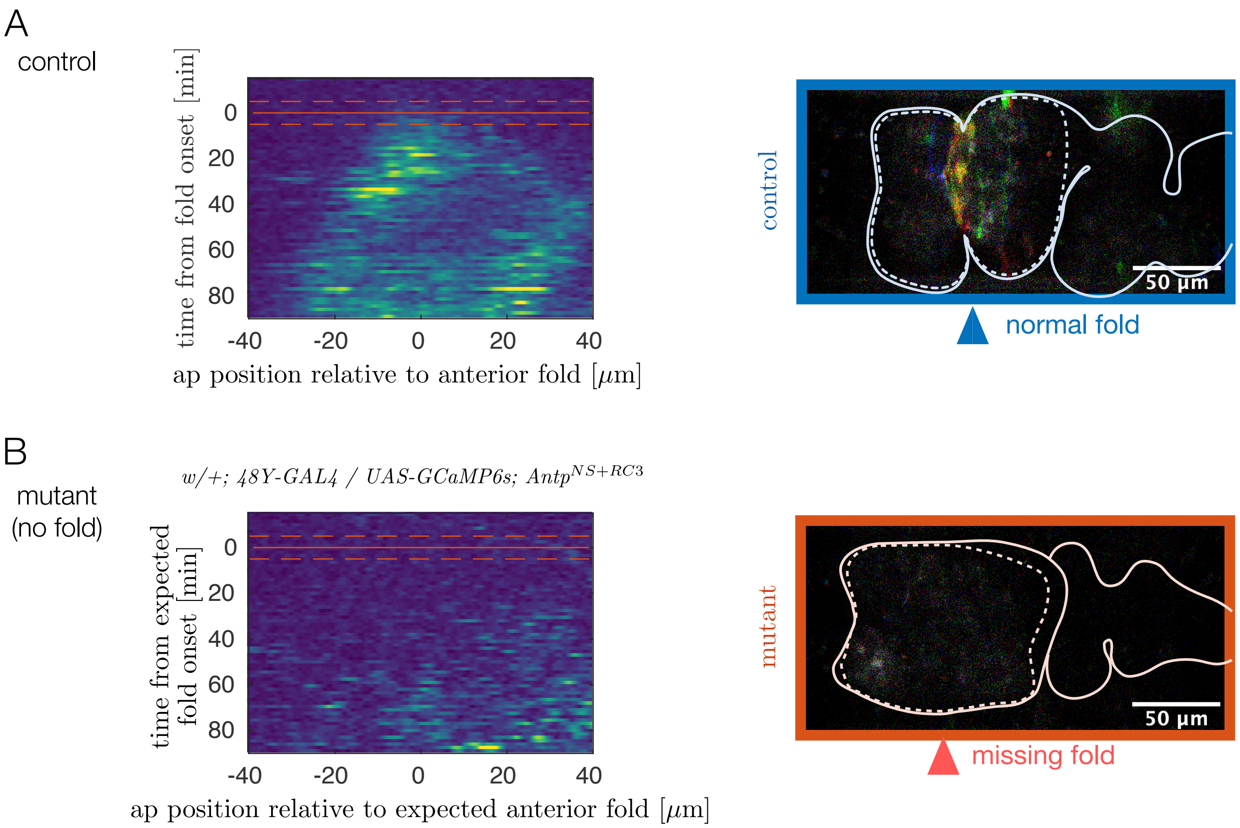


Supplementary Figure S19. ***Antp* mutants show reduced calcium activity in the anterior two chambers for over an hour.** *(A)* Kymographs of wild-type calcium dynamics near the anterior fold for $N = 13$ control embryos show fluctuating calcium activity beginning at the onset of folding at the site of the fold. The domain of calcium activity broadens anteriorly and posteriorly in time. Red solid line marks the onset of the anterior constriction, and dashed lines denote the precision with which this time is known. *(B)* In mutants, almost no calcium pulses are observed during the same timespan. A kymograph of average fluctuating *GCaMP6s* intensity for $N = 5$ *Antp* mutants remains quiescent (dark blue). Given that the anterior fold does not form, $t = 0$ was prescribed based on the depth of the posterior fold. The expected position of the anterior fold (which defines the horizontal axis of the kymograph) is inferred from the position of the anterior fold relative to the anterior face of the midgut in control embryos.

that the result is not sensitive to the choice of analysis. For example, we also computed the mean number of folds formed for each condition and compare the two distributions, as shown in Fig. S20A and B. The mean number of folds formed was reduced in both *Mef2-GAL4* × *UAS-SERCA.R751Q* embryos and *tub67-GAL4; tub16-GAL4* × *UAS-MLCK RNAi* embryos ($p = 3 \times 10^{-8}$ and $p = 0.002$, respectively).

---


[1] C. Englund, A. Birve, L. Falileeva, C. Grabbe, and R. H. Palmer, Development Genes and Evolution **216**, 10 (2006).
[2] D. L. Garaulet, D. Foronda, M. Calleja, and E. Sánchez-Herrero, Development **135**, 3219 (2008).
[3] M. A. Crickmore, V. Ranade, and R. S. Mann, PLOS Genetics **5**, e1000633 (2009).
[4] U. Krzic, S. Gunther, T. E. Saunders, S. J. Streichan, and L. Hufnagel, Nature Methods **9**, 730 (2012).
[5] S. J. Streichan, M. F. Lefebvre, N. Noll, E. F. Wieschaus, and B. I. Shraiman, eLife **7**, e27454 (2018).
[6] S. Preibisch, F. Amat, E. Stamataki, M. Sarov, R. H. Singer, E. Myers, and P. Tomancak, Nature Methods **11**, 645 (2014).
[7] C. B. Phelps and A. H. Brand, Methods **14**, 367 (1998).
[8] N. P. Mitchell and D. J. Cislo, "TubULAR: Tube-like sUrface Lagrangian Analysis Resource for measuring dynamic data of tubular surfaces," (in prep.).
[9] P. Márquez-Neila, L. Baumela, and L. Alvarez, IEEE Transactions on Pattern Analysis and Machine Intelligence **36**, 2 (2014).
[10] T. Chan and L. Vese, IEEE Transactions on Image Processing **10**, 266 (2001).

[11] S. Berg, D. Kutra, T. Kroeger, C. N. Straehle, B. X. Kausler, C. Haubold, M. Schiegg, J. Ales, T. Beier, M. Rudy, K. Eren, J. I. Cervantes, B. Xu, F. Beuttenmueller, A. Wolny, C. Zhang, U. Koethe, F. A. Hamprecht, and A. Kreshuk, Nature Methods **16**, 1226 (2019).
[12] T. G. D. Team, "GIMP," (2019).
[13] W. Thielicke and R. Sonntag, Journal of Open Research Software **9**, 12 (2021).
[14] W. Thielicke and E. Stamhuis, Journal of Open Research Software **2**, e30 (2014).
[15] M. Arroyo and A. DeSimone, Physical Review E **79**, 031915 (2009).
[16] G. Guglielmi, J. D. Barry, W. Huber, and S. De Renzis, Developmental Cell **35**, 646 (2015).
[17] E. Izquierdo, T. Quinkler, and S. De Renzis, Nature Communications **9**, 2366 (2018).

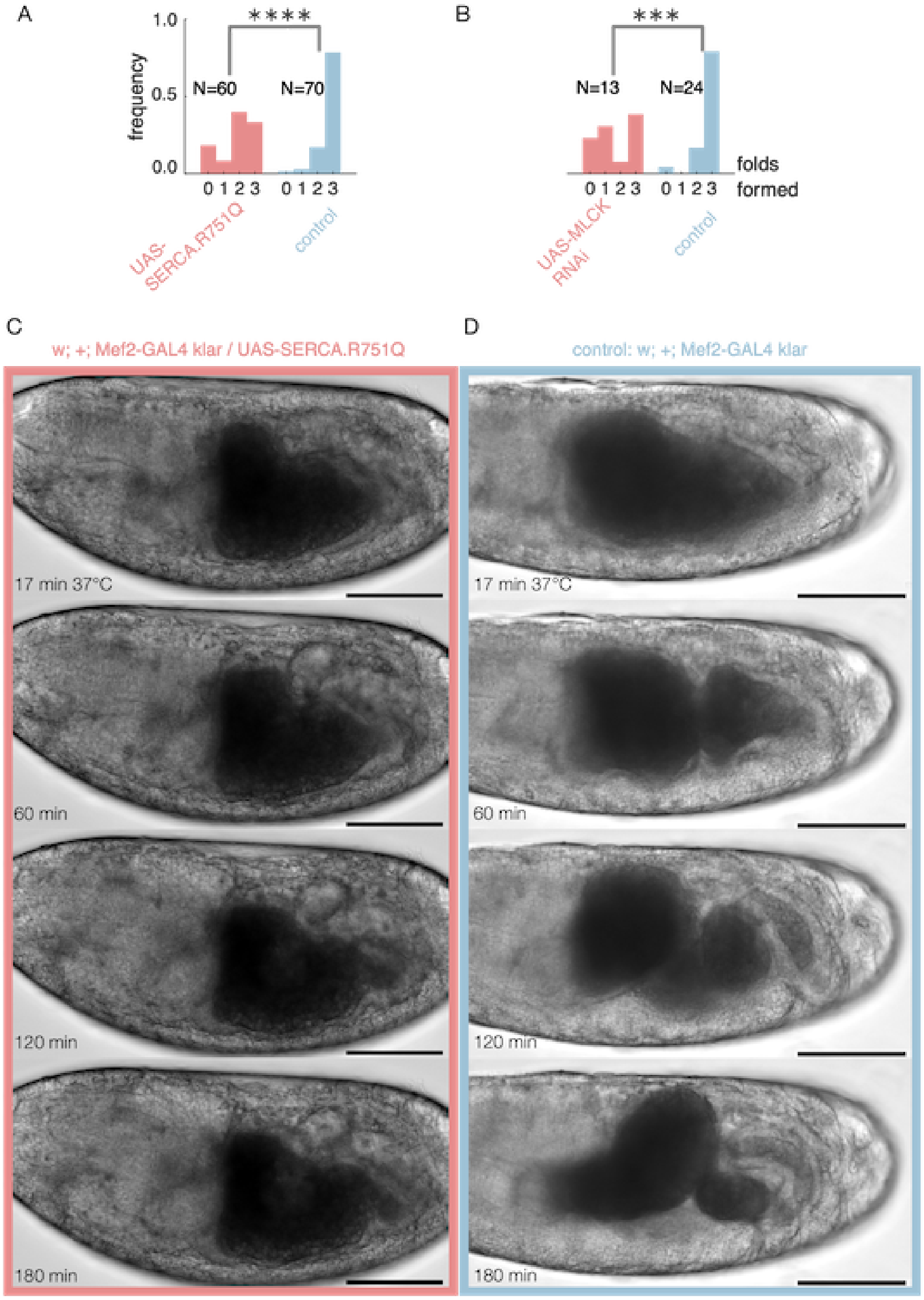


Supplementary Figure S20. **Disrupting normal calcium functioning hinders fold formation.** *(a)* Embryos expressing a dominant negative form of SERCA have fewer successful folds on average (single tailed Z-test, $N = 130$, $p = 3 \times 10^{-8}$) using a muscle-specific driver, *Mef2-GAL4*, at elevated temperatures (continuous heatshock at 37°C). *(b)* Embryos expressing RNAi against MLCK have fewer successful folds on average (single tailed Z-test, $N = 37$, $p = 0.002$). Here we use a ubiquitous driver, *tub15-GAL4; tub67-GAL4* at continuous heatshock at 37°C. *(c)* Brightfield imaging of embryos expressing a mutant form of *SERCA* in muscles show reduced folding activity. Here, driving a mutant *SERCA* expression via heatshock starting at stage 15a shows no folds. *(d)* Control embryos without the mutant form of *SERCA*, in contrast, typically form three folds. Timestamps denote minutes since the onset of heatshock and scalebar is 100 $\mu$m.
